# Jagged1 regulates extracellular matrix deposition and remodeling in triple-negative breast cancer

**DOI:** 10.1101/2025.05.23.655829

**Authors:** Marjaana Parikainen, Ujjwal Suwal, Pekka Rappu, Jyrki Heino, Cecilia Sahlgren

## Abstract

The extracellular matrix (ECM) in the tumor microenvironment plays an essential role in cancer progression. High expression of Jagged1 correlates with poor patient survival and promotes *in vivo* tumor growth and invasion in triple-negative breast cancer (TNBC). Using transcriptomics, proteomics, and imaging of cancer cell/fibroblast co-cultures, both *in vitro* and *in vivo*, we demonstrate that Jagged1-mediated crosstalk between TNBC cells and fibroblasts enhances myofibroblast activation, collagen deposition, and alignment of ECM fibers. Jagged1 increases transforming growth factor beta (TGFβ) activity in fibroblast co-cultures and the Jagged1-induced ECM alignment can be prevented by a TGFβ inhibitor. Thus, Jagged1 regulates ECM remodeling upstream of TGFβ. Furthermore, higher substrate stiffness increases Jagged1 expression, pointing towards a feed-forward loop between Jagged1, ECM stiffness, and TGFβ. With the emergence of safe therapeutics targeting specific Notch ligands and receptors, Jagged1 modulation may offer a path to therapeutically target invasive breast cancer.

## Main

Tumor progression and metastasis depend on the cancer cell-intrinsic changes and the formation of a supportive tumor microenvironment (TME)^1,2^. The TME consists of various types of non-malignant cells, such as immune cells, endothelial cells, adipocytes, and fibroblasts, surrounded by extracellular matrix (ECM). The ECM is a complex and dynamic network of cross-linked proteins that coordinates cellular behavior in multicellular organisms. Increased matrix deposition, stiffness, and a more aligned ECM organization, stimulated by such key signaling pathways as TGFβ and hypoxia signaling, drive tumor cell malignancy and metastasis^3–5^. The stiff ECM of tumors is increasingly recognized as a major factor hindering immune infiltration and promoting therapy resistance, and thus understanding the mechanisms driving ECM dysregulation in the TME is of utmost importance for the development of novel cancer therapeutics.

In malignant tumors, cancer-associated fibroblasts (CAFs) play a critical role in synthesis and organization of the ECM. Diverse cellular origin, spatial location, and inherent plasticity of the CAFs make them a versatile cell type for which three major phenotypes have been described: myofibroblastic CAFs (myCAFs), inflammatory and growth factor-enriched CAFs (iCAFs), and antigen-presenting CAFs (apCAFs)^6^. myCAFs actively secrete ECM proteins, such as fibronectin and collagens, and ECM modifying enzymes, including matrix metalloproteases (MMPs) and lysyl oxidase (LOX)^6–9^. myCAFs are typically contractile and express α-smooth muscle actin (αSMA). In different tumors, myCAFs may either prevent or promote cancer progression. Breast cancer is the leading cause of cancer death in women, complicated by molecular subtypes defined by estrogen receptor (ER), progesterone receptor (PR), and human epidermal growth factor receptor 2 (HER2) status. Triple-negative breast cancer (ER-/PR-/HER2-) is aggressive and treatment-resistant, underscoring the need for new therapies. In breast cancer, the abundance of αSMA positive myCAFs correlates with a more aggressive phenotype and disease relapse^10,11^. Multiple paracrine factors within the TME have been reported to induce the differentiation of stromal fibroblasts or iCAFs into myCAFs, most importantly TGFβ. Emerging evidence suggests that juxtacrine cell–cell contact-mediated mechanisms may also play a role in this context^12–17^.

The Notch signaling pathway has a critical role in embryogenesis and maintenance of most tissues, and aberrant Notch signaling is frequently observed in various cancers^18^. Notch signaling is activated when a Notch ligand interacts with a Notch receptor in a cell–cell contact-dependent manner. Upon activation, the receptor undergoes two subsequent cleavages to release its intracellular domain, which then translocates to the nucleus to activate target gene expression. Mammalian cells express four Notch receptors (Notch1-4) and five Notch ligands (Jagged1/2, Dll1/3/4). Notch activating cell–cell contacts in cancer may occur either between cancer cells or cancer cells and cells of the TME. Both oncogenic and tumor-suppressive effects of Notch have been shown, depending on the context, and Notch signaling is involved in all hallmarks of cancer^19^.

In breast cancer, the Notch signaling pathway has been implicated in tumor progression and TNBC^20^. Most of the research has been focusing on the role of the Notch receptors in breast cancer pathogenesis, while comprehensive studies on the functional roles of Notch ligands are lagging behind. The Notch ligand Jagged1 is overexpressed in several cancer types and participates in many tumor-promoting activities, such as increased proliferation, vascularization, and invasion^21^. In breast cancer, high Jagged1 expression has been associated with the aggressive basal subtype and an increased risk of metastasis^22–26^, but a more detailed understanding of Jagged1-mediated tumorigenic functions is still missing.

In this study, we show that Jagged1 promotes the *in vivo* growth and invasion of TNBC cells. High expression of Jagged1 predicts poor patient survival, specifically in aggressive breast cancer subtypes (HER2-enriched and basal-like/TNBC). Using previously published single-cell RNA sequencing (scRNA-seq) data^27^, publicly available breast cancer patient data^28^, and a 3D *in vitro* model, we demonstrate that Jagged1 enhances invasiveness, cancer stem cell-like (CSC) features, and expression of ECM-related genes in TNBC cells. High Jagged1 in cancer cells correlates with an increased differentiation of myCAFs with a Notch and TGFβ gene signature in scRNA-seq data of TNBC tumors. Interestingly, when Jagged1-expressing cancer cells were co-cultured with fibroblasts, collagen production and ECM fiber alignment were highly increased in a TGFβ-dependent manner. Furthermore, increasing substrate stiffness elevates Jagged1 expression levels in cancer cells, pointing towards a feed-forward loop between tumor-promoting ECM remodeling and Jagged1. Finally, we validate the discovered relationship between Jagged1, TGFβ, myCAF activation, and increased collagen deposition utilizing *in vivo* co-culture tumors and breast cancer patient datasets. Taken together, our data designate Jagged1 as a central regulator of TGFβ activity and ECM remodeling in TNBC, thus promoting cancer progression.

## Results

Jagged1 enhances the aggressive behavior of triple-negative breast cancer cells *in vivo* To investigate how Jagged1 affects breast cancer progression, we first assessed Jagged1 expression levels in scRNA-seq data^27^ of cancer cells from different subtypes of breast cancer. Jagged1 was more highly expressed in the aggressive breast cancer subtypes (HER2-enriched and TNBC) while expressed at a low level in ER+ cancer cells (Fig. 1a-b). Interestingly, by utilizing published patient datasets^29^, we found that high Jagged1 expression was associated with worse relapse-free survival in the same aggressive subtypes where it is more highly expressed (Fig. 1c). High Jagged1 expression has been previously linked to ER-breast cancer^23,30,31^. We also confirmed that Jagged1 was highly expressed in a panel of ER-breast cancer cell lines and low in ER+ cell lines (Extended Data Fig. 1a). High Jagged1 expression led to worse survival in a cohort of patients with ER-breast cancer while having the opposite effect in ER+ breast cancer (Extended Data Fig. 1b), further supporting the notion that Jagged1 drives breast cancer progression in the absence of estrogen stimulation. To clarify the role of Jagged1 in aggressive breast cancer, we generated a Jag1 CRISPR-Cas9 knockout cell line out of MDA-MB-231 human TNBC cells (Jag1KO). First, we assessed the ability of Jag1KO cells to form tumors *in vivo* in the chick chorioallantoic membrane (CAM) model in comparison to the parental MDA-MB-231 wild-type cell line that retains a high Jagged1 expression (Jag1WT). Loss of Jagged1 led to a significant decrease in tumor growth (Fig. 1d). In addition, when injected intravenously into zebrafish embryos, Jag1KO cells showed decreased survival in the circulation and reduced formation of metastases (Fig. 1e-g). Altogether, our data show that Jagged1 increases TNBC cell growth and invasion *in vivo* and is associated with poor prognosis in aggressive, ER-breast cancer.

**Figure 1.**
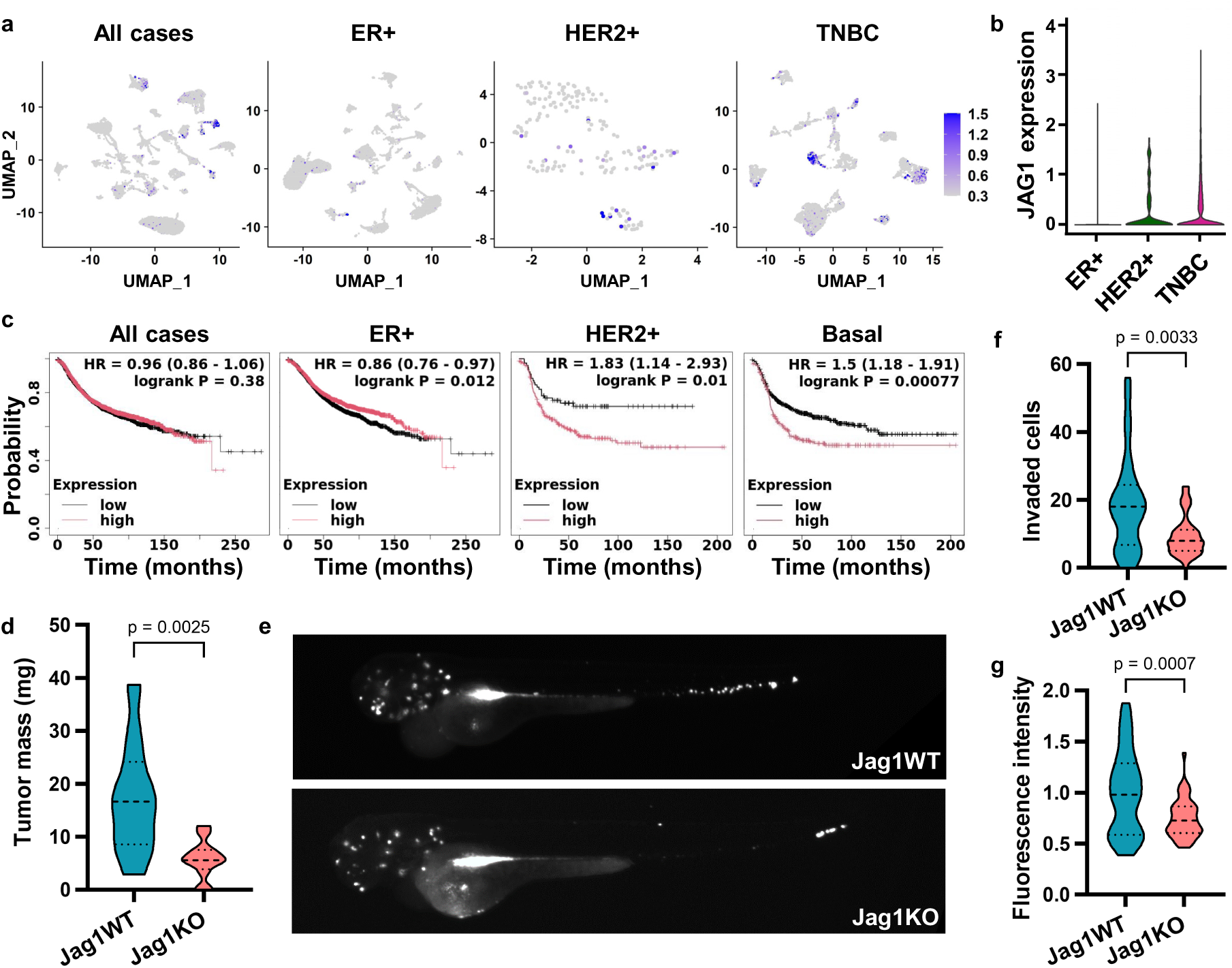
Jagged1 promotes breast cancer aggressiveness. **a,b** *JAG1* expression in scRNA-seq data of cancer cells from different subtypes of breast cancer. **c** Relapse-free survival of patients with low (black) or high (red) expression of Jag1 in different subtypes of breast cancer. **d** Tumor masses of MDA-MB-231 wild-type (Jag1WT) and Jag1 knockout (Jag1KO) cell chick chorioallantoic membrane xenograft models. p-values are calculated by two-tailed unpaired t-test. n = 10 tumors for Jag1WT and n = 12 for Jag1KO. **e** Representative images of zebrafish embryos injected intravenously with fluorescently labeled cancer cells. **f** Quantification of cells that have invaded into the tails of zebrafish embryos. p-value is calculated by two-tailed unpaired Mann-Whitney test. n = 30 embryos for Jag1WT and n = 42 for Jag1KO. **g** Normalized fluorescence intensity of cells invaded to the heads of zebrafish embryos. p-value is calculated by two-tailed unpaired t-test. n = 30 embryos for Jag1WT and n = 44 for Jag1KO.

### Jagged1 regulates TNBC cell phenotype and gene expression

To clarify the mechanisms behind Jagged1-mediated TNBC progression, we split cancer cells of the scRNA-seq dataset^27^ into cells with either high (Jag1 high) or low (Jag1 low) expression of *JAG1*. TNBC tumors had the highest percentage of Jag1 high cells (Fig. 2a-b). A gene set enrichment analysis (GSEA) of genes differentially expressed between Jag1 high and Jag1 low cells (Supplementary Table 1) revealed ECM-related genes as the most significantly upregulated genes in Jag1 high cells (Fig. 2c). In addition, Jag1 high cells displayed increased expression of genes involved in the Notch, Wnt, and TGFβ pathways, along with increased expression of breast CSC and epithelial-mesenchymal transition (EMT) markers (Fig. 2c-f). We set up 3D Matrigel cultures of MDA-MB-231 cells to assess the impact of Jagged on growth and invasiveness and elucidate the Jagged-driven transcriptome. After 7 days of culture, Jag1KO spheroids were less invasive and smaller compared to Jag1WT spheroids and appeared more differentiated (Fig. 2g). A similar phenotype was seen with siRNA-mediated silencing of Jagged1 while reintroducing Jagged1 into Jag1KO cells saved the phenotype (Extended Data Fig. 2a). The effect seemed to be Notch signaling dependent, as inhibition of Notch by using a gamma-secretase inhibitor or knocking out Notch1 induced a similar phenotype (Extended Data Fig. 2b). A genome-wide transcriptome analysis on these spheroid samples revealed hundreds of genes differentially expressed depending on the Jagged1 expression status (Fig. 2b, Supplementary Table 2). We validated the RNA-sequencing results of a few hit genes by quantitative PCR (Extended Data Fig. 2c). A GSEA showed that ECM and cell migration-related gene ontology (GO) terms were significantly over-represented among genes upregulated in Jag1WT cells (Fig. 2i). Several collagens, integrins, and matrix remodeling enzymes were found among the Jagged1-regulated ECM genes (Supplementary Table 2). Notably, highly similar gene sets were correlated with Jagged1 expression level in a human breast cancer patient database (Fig. 2j, Supplementary Table 3). In conclusion, higher Jagged1 expression correlated with increased CSC-like features and invasiveness both *in vitro* and *in vivo* in breast cancer patient data.

**Figure 2.**
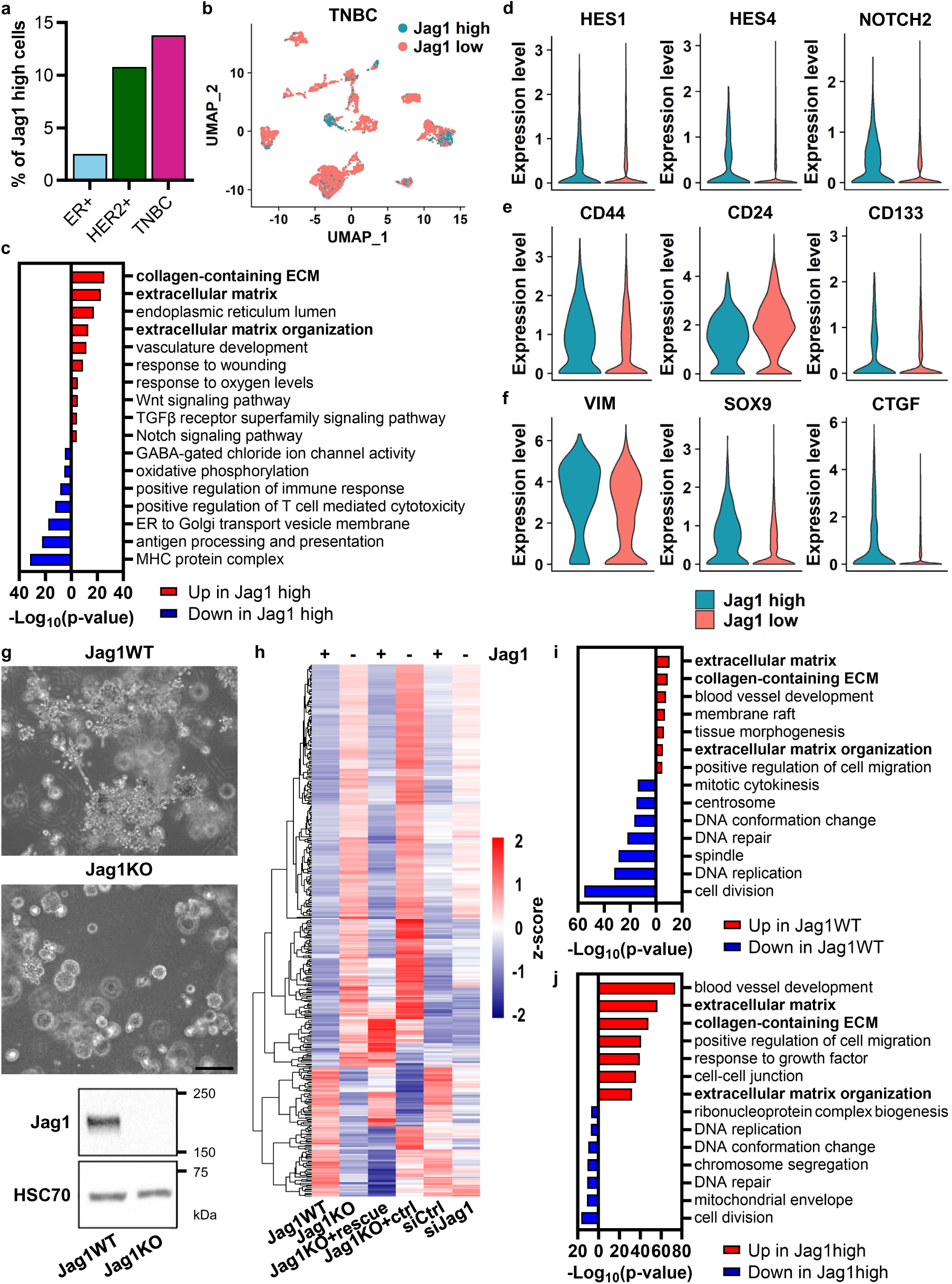
Jagged1 enhances invasiveness, cancer stem cell-like features, and expression of ECM genes in TNBC cells. **a** Percentage of cancer cells with high *JAG1* expression in scRNA-seq data of different breast cancer subtypes. **b** Uniform manifold approximation projection (UMAP) visualization of TNBC cells split into Jag1 high and Jag1 low populations based on *JAG1* expression level. **c** Gene set enrichment analysis (GSEA) of genes differentially expressed between Jag1 high and Jag1 low TNBC cells in scRNA-seq data (fdr < 0.01 and average log2 fold change > 0.500). Examples of **d** Notch pathway genes, **e** breast cancer stem cell markers, and **f** epithelial-mesenchymal transition markers differentially expressed between Jag1 high and Jag1 low TNBC cells in scRNA-seq data. n = 719 of Jag1 high and n = 4494 Jag1 low TNBC cells. **g** Representative images of MDA-MB-231 Jag1WT and Jag1KO 3D spheroids and a western blot showing Jag1 expression in the cell lines. Scale bar 200 µm. **h** A clustered heatmap of a genome-wide transcriptome analysis, showing z-scores of genes differentially expressed between spheroid samples of Jag1WT and Jag1KO cells, Jag1KO cells transfected with Jag1-containing plasmid (Jag1KO+rescue) or empty vector plasmid (Jag1KO+ctrl), and Jag1WT cells transfected with non-targeting siRNA (siCtrl) or Jag1-targeting siRNA (siJag1) (fdr < 0.05 and fold change > 1.5). **i** GSEA of genes differentially expressed between Jag1WT and Jag1KO spheroids (fdr < 0.05). **j** GSEA of genes either positively (red) or negatively (blue) correlating with *JAG1* expression (Spearman’s correlation coefficient >0.300 or <-0.300, respectively) in a breast cancer patient mRNA expression dataset (TCGA, Cell 2015, n = 817).

### Jagged1 induces ECM deposition and remodeling in fibroblast co-cultures via TGFβ signaling

ECM components can, in principle, be produced by any cell, and it has been shown that cancer cells can have an altered expression of for example different collagens and matrix remodeling enzymes^29,30^. However, the major ECM producers are fibroblasts, and especially in the context of cancer, the myCAFs. Since Jagged1 upregulated the expression of ECM-related genes in cancer cells, we assessed how Jagged1 expressed by cancer cells affects ECM production by fibroblasts. For this, we first checked the abundance of Jag1 high cancer cells in individual TNBC patients in the scRNA-seq data and then split patients into Jag1 high or Jag1 low groups based on whether they had higher or lower percentage of Jag1 high cells than the average (Fig. 3a-b). We then performed a GSEA of genes differentially expressed in fibroblasts from Jag1 high and Jag1 low patients (Supplementary Table 4). Fibroblasts from tumors with high *JAG1* expression in cancer cells showed increased expression of *NOTCH3* and Notch target genes, *TGFB1* and other TGFβ pathway genes, as well as myCAF and cellular contractility markers (Fig. 3c-f), while the expression of several iCAF markers was downregulated (Extended Data Fig. 3a).

**Figure 3.**
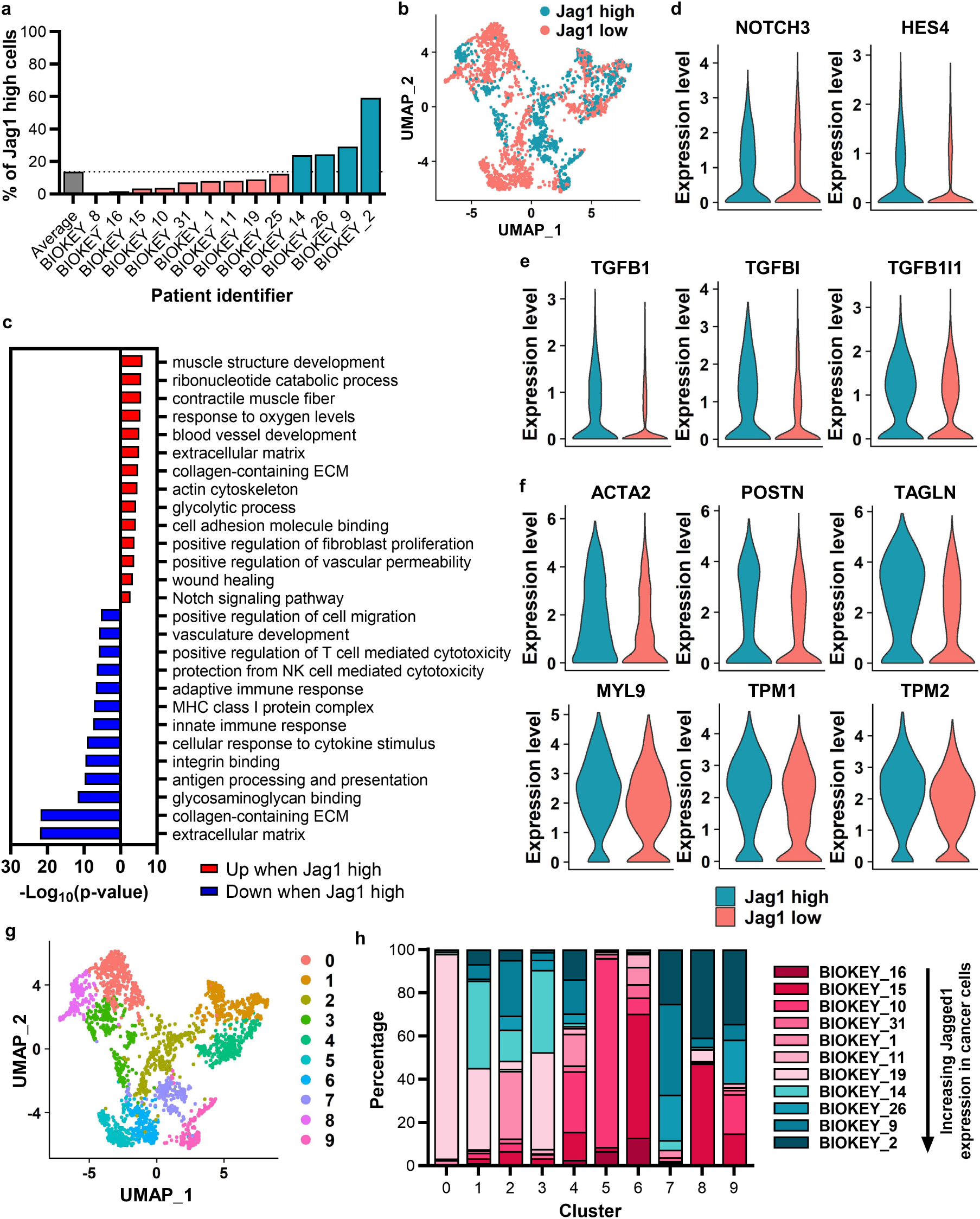
High Jagged1 in cancer cells promotes Notch and TGFβ pathways and cellular contractility in fibroblasts. **a** Percentage of cancer cells with high *JAG1* expression in scRNA-seq data of TNBC samples from individual patients. **b** UMAP visualization of fibroblasts from patients with either high or low Jag1 in cancer cells. **c** GSEA of genes differentially expressed between fibroblasts from patients with high or low Jag1 in cancer cells (fdr < 0.01 and average log2 fold change > 0.500). Expression of **d** Notch pathway genes, **e** TGFβ pathway genes, and **f** myofibroblast markers in fibroblasts from patients with high or low Jag1 in cancer cells. **g** UMAP visualization of fibroblast clusters. **h** Percentage of fibroblasts from each individual patient in fibroblast clusters. n = 881 of Jag1 high fibroblasts and n = 1530 of Jag1 low fibroblasts.

In contrast to the upregulation of ECM-related genes observed in Jag1 high cancer cells, there was a Jagged1-dependent shift in the composition of the expressed ECM genes in fibroblasts, with some genes being upregulated and others downregulated (Extended Data Fig. 3b-c). There were no significant differences in the expression of fibrillar collagens, while several TGFβ-regulated collagen fiber modifying enzymes were upregulated in Jag1 high fibroblasts (Extended Data Fig. 3d). As we saw an increase in the expression of myCAF markers and a decrease in iCAF markers in Jag1 high fibroblasts, we hypothesized that Jagged1 induces a phenotypic switch of the fibroblasts and thus affects their clustering in the scRNA-seq data. When we clustered the fibroblasts, we noticed one cluster consisting almost entirely of fibroblasts from the tumors with high Jagged1 in cancer cells (Fig. 3g-h). This fibroblast cluster had highly upregulated Notch and TGFβ pathway signatures, while also demonstrating upregulated expression of myCAF markers, fibrillar collagens, and ECM modifying enzymes (Extended Data Fig. 4a-e, Supplementary Table 5). Interestingly, the gene markers of this cluster showed high similarity to the markers of the specific myCAF subpopulation, TGFβ-myCAFs, previously discovered by Kieffer et al.^31^, while *JAG1* expression correlated significantly with the expression of TGFβ-myCAF markers in breast cancer patient data (Extended Data Fig. 3f). This suggests that Jagged1 induces activation of this specific myCAF subpopulation in breast cancer.

As the Jag1 low tumors in the scRNA-seq data contained some cancer cells expressing high levels of Jagged1, we cannot rule out that some of the Jag1 low fibroblasts may still have been activated by Jagged1. To clarify the effect of Jagged1 on fibroblasts and ECM, we co-cultured our Jag1WT or Jag1KO cells with mouse embryonic fibroblasts (MEFs) as spheroids in a U-bottom well 3D culture system without any external added ECM. A mass spectrometry analysis on matrisome-enriched protein samples from these spheroids revealed a significant upregulation of collagen production in Jag1WT co-cultures (Fig. 4a). The use of mouse fibroblasts and human cancer cells allowed us to perform a species-specific analysis on the proteomics results, showing cell type-specific changes in protein expression (Fig. 4b). Upregulation of collagen production occurred both in cancer cells and in fibroblasts. However, when we stained spheroids with the fluorescent pan-collagen probe CNA35^32^, collagen mostly accumulated in the areas where fibroblasts resided in the spheroids (Fig. 4c). Another interesting observation was an apparent dysregulation of the TGFβ pathway, as several proteins related to the pathway exhibited Jagged1-dependent expression (Fig. 4b). Previously we had seen a Jagged1-dependent upregulation of a TGFβ gene signature both in cancer cells and in fibroblasts in the scRNA-seq data (Fig. 2c and Fig. 3e). Western blot analysis of spheroid samples showed increased phosphorylation of both SMAD2 and SMAD3, indicating increased TGFβ activity in the Jag1WT co-cultures (Fig. 4d). Collagen production was increased by almost 50 % in the presence of Jagged1, whereas there was no difference in the production of fibronectin. The differences in TGFβ activity and collagen production were even more striking when cancer cells were co-cultured with human telomerase immortalized fibroblasts (TIFs), while fibronectin appeared to be downregulated in Jag1WT+TIF co-cultures (Fig. 4e). The myCAF marker αSMA was also upregulated by Jagged1 in both fibroblasts. Altogether, these results suggest that Jagged1 expressed by cancer cells induces collagen production and myCAF activation via increased TGFβ activity.

**Figure 4.**
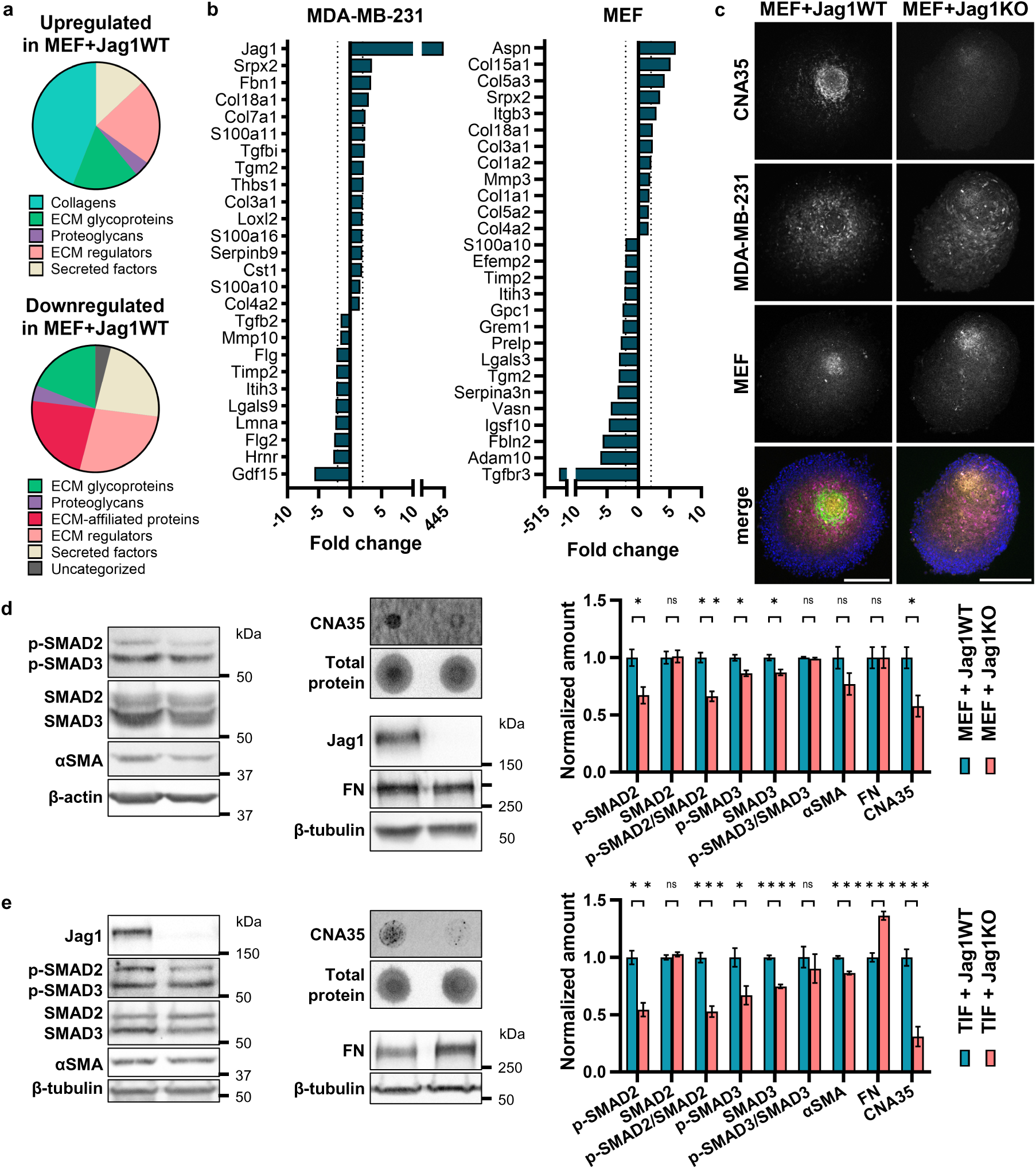
Jagged1 signaling from cancer cells induces fibroblast activation, collagen production, and TGFβ signaling in 3D co-cultures. **a** MatrisomeDB classifications of proteins either upregulated or downregulated in MDA-MB-231 Jag1WT and mouse embryonic fibroblast (MEF) co-culture spheroids compared to Jag1KO co-cultures. **b** Proteins differentially expressed by either MDA-MB-231 cells or MEFs in MEF+Jag1WT co-cultures compared to MEF+Jag1KO co-cultures. **c** Cross sections of MDA-MB-231 (magenta) and MEF (orange) co-culture spheroids stained with CNA35 collagen probe (green) and DAPI (blue). Scale bar 200 µm. Western blots and dot blots of indicated proteins in co-culture spheroids of MDA-MB-231 cells and **d** MEFs or **e** human telomerase immortalized fibroblasts (TIF). Quantifications of four independent experiments are shown to the right. Data are presented as mean ± SEM. p-values are calculated by two-tailed unpaired t-test. CNA35, collagen-binding adhesion protein tagged with EGFP. FN, fibronectin. *p<0.05, **p<0.01, ***p<0.001, ****p<0.0001.

Along with the increased deposition of ECM, especially fibrillar collagens, the increased cross-linking and stiffness of the ECM lead to the formation of a fibrotic, tumor-promoting microenvironment. ECM surrounding the tumor is often anisotropically organized, and the densely packed and highly aligned ECM fibers create tracks for efficient cancer cell migration, thus promoting metastasis^33,34^. As we already established a significant increase in collagen deposition and altered expression of several matrix remodeling factors upon Jagged1 stimulation, we wanted to evaluate whether Jagged1 is involved in the regulation of ECM fiber organization. Interestingly, when fibroblasts were co-cultured with Jagged1-expressing cancer cells, the fibronectin and collagen fibers of cell-derived matrices (CDMs) were highly aligned compared to fibroblast monocultures or co-cultures with Jag1KO cells (Fig. 5a-c and Extended Data Fig. 5a-c). We hypothesized this effect might be TGFβ-dependent, as Jagged1 was shown to induce TGFβ activity. Indeed, inhibition of TGFβ prevented the Jagged1-induced alignment of ECM fibers completely (Fig. 5d-g), whereas recombinant TGFβ1 treatment rescued the Jag1KO phenotype (Fig. 5h-k). Similar Jagged1-dependent ECM alignment was observed in another TNBC cell line, MDA-MB-436 (Extended Data Fig. 6a-c). The induction of fiber alignment was dependent on Notch activation, as the gamma-secretase inhibitor PF-03084014 prevented the alignment of Jag1WT CDMs (Extended Data Fig. 6d-e). Intriguingly, recombinant Jagged1 coating on the coverslips was able to induce an increase in fibroblast-produced CDM fiber alignment, whereas Notch ligands Dll1 and Dll4 lacked this ability (Extended Data Fig. 7). Both the transcriptomics and the proteomics analyses of Jag1WT samples revealed an upregulation of matrix crosslinking lysyl oxidases, and inhibition of LOX also partly prevented the Jagged1-induced fiber alignment, although not as efficiently as Notch or TGFβ inhibition (Extended Data Fig. 8). Expression of Jagged1 was increased upon culturing MDA-MB-231 as well as the ER+ MCF7 WT cells on increasing substrate stiffness, pointing towards a general feed-forward loop between ECM stiffness and Jagged1 expression, as both increased deposition and higher alignment of matrix fibers are indicative of increased tissue stiffness (Extended Data Fig. 9). Taken together, Jagged1 is a central regulator of both ECM deposition and remodeling, inducing matrix fiber alignment through Notch and TGFβ signaling activity.

**Figure 5.**
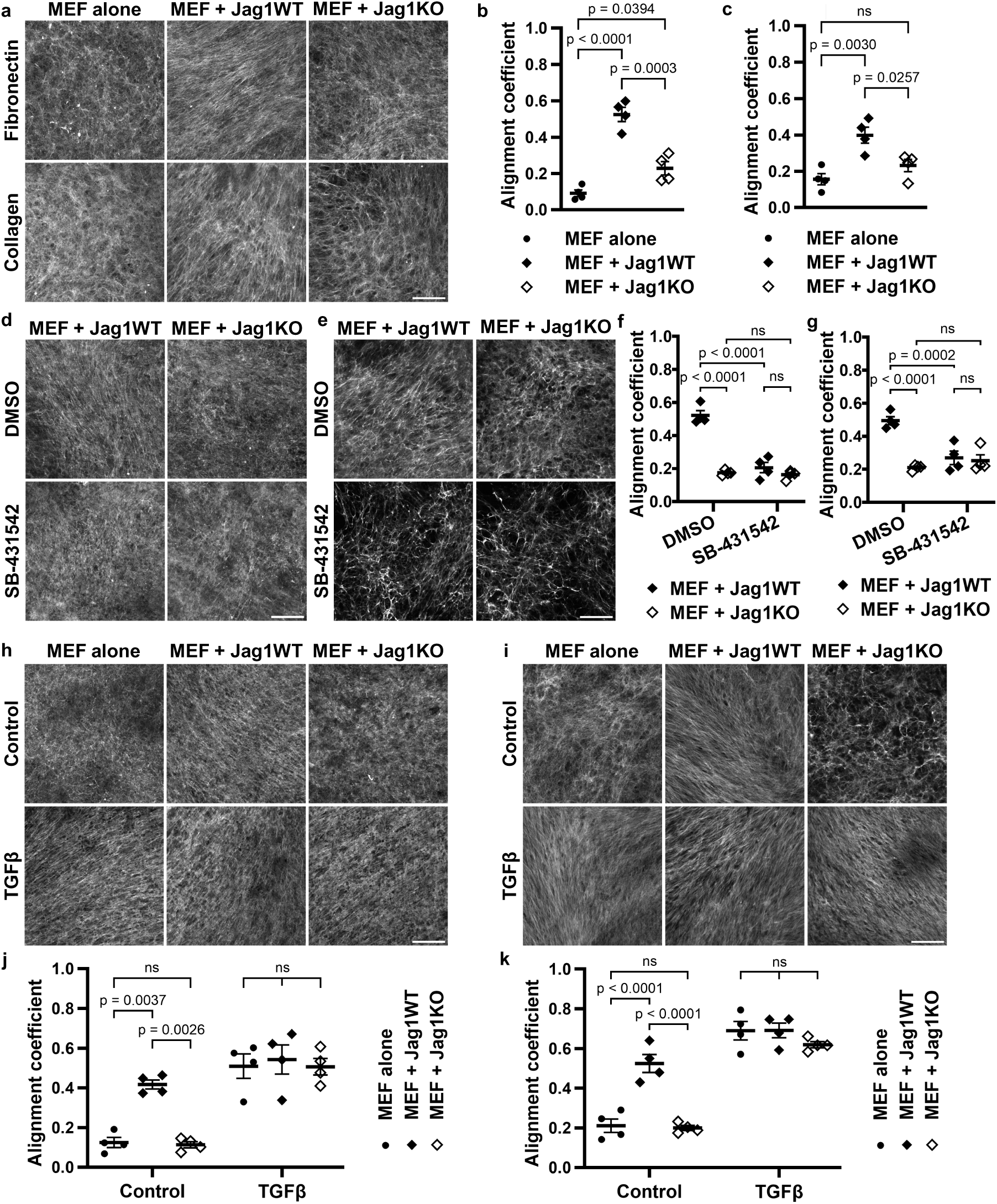
Jagged1 directs extracellular matrix fiber alignment via TGFβ signaling. **a** Representative images of cell-derived matrices (CDM) from MEF monocultures or MEF co-cultures with MDA-MB-231 Jag1WT or Jag1KO cells, and alignment coefficients of **b** fibronectin and **c** collagen fibers. p-values are calculated by one-way ANOVA with Tukey’s multiple comparisons test. Representative images and fiber alignment coefficients of **d,f** fibronectin and **e,g** collagen of CDMs from DMSO or SB-431542 TGFβ inhibitor treated Jag1WT or Jag1KO co-cultures with MEF cells. p-values are calculated by two-way ANOVA. Representative images and fiber alignment coefficients of **h,j** fibronectin and **i,k** collagen of CDMs from control or recombinant TGFβ1 treated MEF monocultures or MEF co-cultures with Jag1WT or Jag1KO cells. p-values are calculated by two-way ANOVA with Šídák’s multiple comparisons test. Data are presented as mean ± SEM of four independent experiments. Scale bar 100 µm.

### Jagged1 promotes fibroblast activation and collagen deposition *in vivo*

To determine if Jagged1 regulates fibroblast behavior and ECM production *in vivo*, we co-cultured MEFs with either Jag1WT or Jag1KO cells in the CAM model. Jag1WT tumors were significantly bigger, and based on hematoxylin and eosin stainings, they showed more histological resemblance to invasive breast cancer than Jag1KO tumors while also demonstrating stronger eosin staining, possibly pointing to increased collagen (Fig. 6a-b). We next analyzed tumor sections for expression of collagen, fibronectin, and the myCAF marker αSMA (Fig. 6c-e). αSMA positive cells appeared to reside in glandular structures within the tumor, surrounded by pan-cytokeratin-positive cancer cells of epithelial origin. Blood vessels also showed bright αSMA staining due to smooth muscle cells in the vessel walls (Fig. 6d). Similar separation of fibroblast and cancer cell location was seen previously in co-culture spheroids (Fig. 4c). We quantified αSMA expression in these glandular structures and it was significantly higher in Jag1WT tumors, indicating Jagged1-induced myCAF differentiation. There was also a significant increase in the amount of collagen in the whole tumor area, even when cells were grafted in collagen-containing Matrigel. No differences were seen in fibronectin deposition, in line with our previous results from MEF co-culture spheroids (Fig. 4d). Importantly, both αSMA (*ACTA2*) and multiple different collagens showed positive RNA expression level correlation with Jagged1 in human breast cancer patient datasets, while the correlation between fibronectin and Jagged1 was much weaker (Fig. 6f and Extended Data Fig. 10a). Jagged1 expression also correlated positively with many matrix remodeling enzymes, components of the TGFβ pathway, and several integrins in the patient dataset^28^ (Extended Data Fig. 10), as also seen in our RNA-sequencing and proteomics data and in the scRNA-seq data^27^. Altogether, our data establish Jagged1 as an important modulator of the tumor tissue structure upstream of TGFβ in TNBC (Fig. 7).

**Figure 6.**
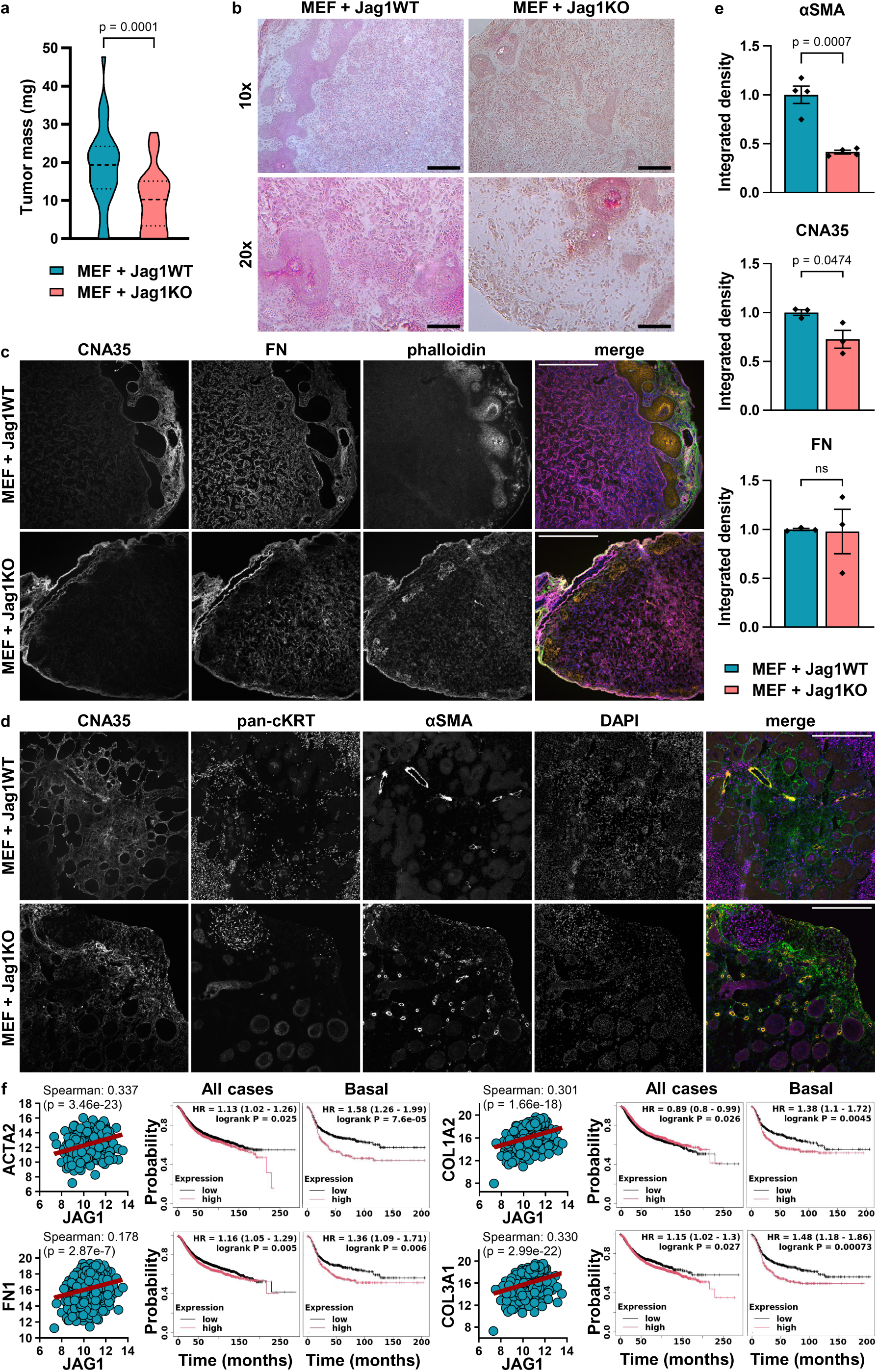
Jagged1 promotes collagen production and αSMA expression *in vivo*. **a** Tumor masses of MDA-MB-231 Jag1WT or Jag1KO cell co-cultures with MEF cells in chick chorioallantoic membrane (CAM) xenograft model. p-values are calculated by two-tailed unpaired t-test. n = 37 tumors for Jag1WT and n = 40 for Jag1KO. **b** H&E stained tissue sections of CAM tumors in **a**. Scale bars 200 µm and 100 µm for 10x and 20x magnifications, respectively. CAM tumor cryosections stained for **c** collagen (CNA35; green), fibronectin (FN; magenta), actin (phalloidin; orange), and nuclei (DAPI; blue), and for **d** collagen (CNA35; green), pan-cytokeratin (pan-cKRT; magenta), αSMA (orange), and nuclei (DAPI; blue). Scale bar 500 µm. **e** Quantifications of αSMA, CNA35 (collagen), and fibronectin (FN) fluorescence integrated densities in CAM tumor cryosections. n = 4 tumors for αSMA and n = 3 tumors for collagen and fibronectin in each group. αSMA was quantified from glandular structures containing fibroblasts, and collagen and fibronectin were quantified from whole tumor area, excluding possible surrounding CAM. Data are presented as mean ± SEM. p-values are calculated by two-tailed unpaired t-test. **f** mRNA expression level correlation with Jag1 mRNA expression in breast cancer patients (TCGA, Cell 2015, n = 817) and relapse-free survival of patients with low (black) or high (red) expression of ACTA2, FN1, COL1A2, or COL3A1 in all breast cancer cases and in basal breast cancer separately.

**Figure 7.**
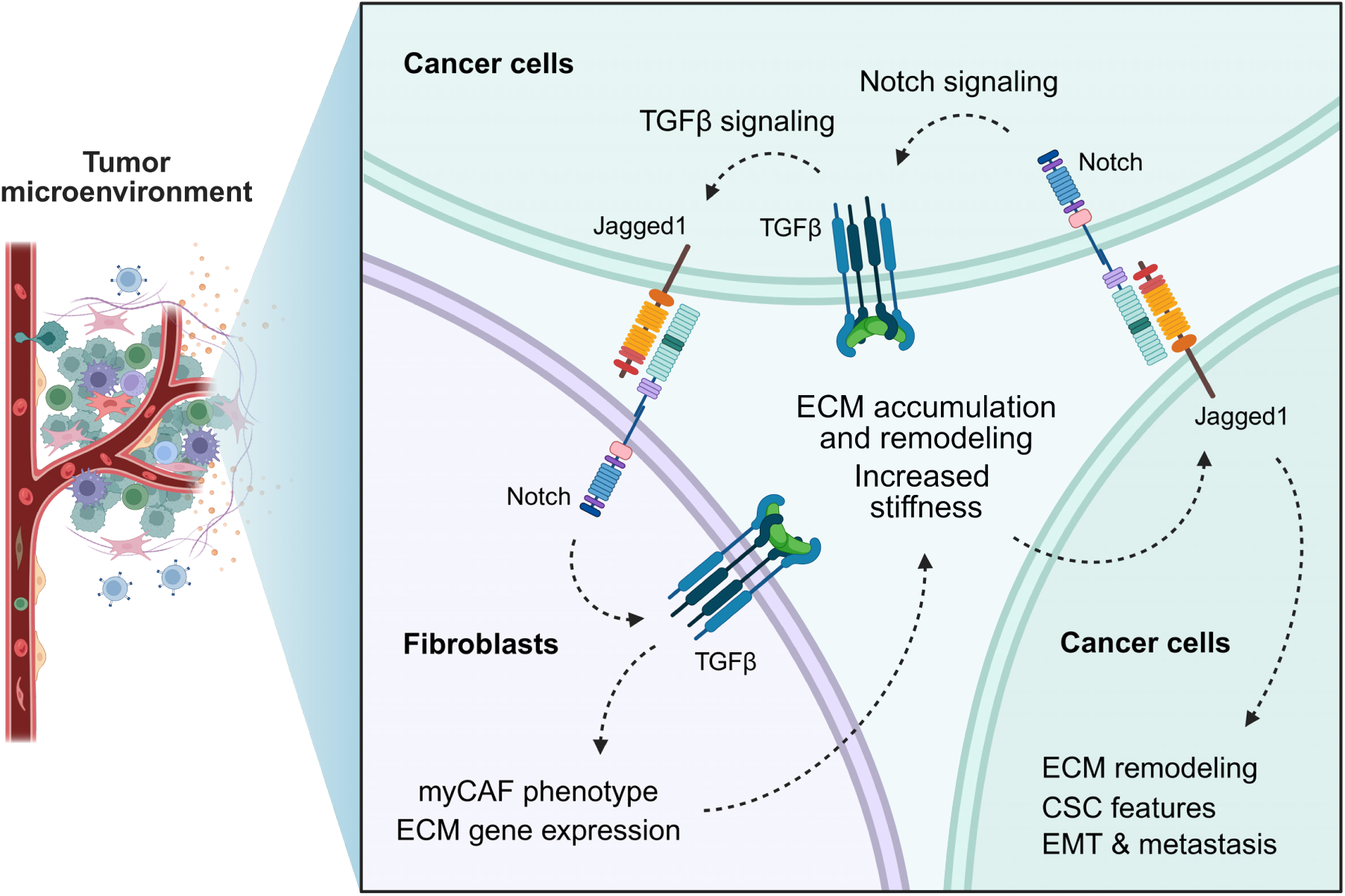
Jagged1-mediated crosstalk between TNBC cells and fibroblasts enhances myofibroblast activation, ECM accumulation, and alignment of ECM fibers via increased Notch activation and TGFβ activity. High Jagged1 expression promotes cancer stem cell-like features, EMT, and invasiveness of TNBC cells, thus enhancing tumor formation and metastasis. Jagged1 expressed by cancer cells activates Notch in the surrounding fibroblasts and increases TGFβ activity. Fibroblasts interacting with Jagged1-expressing cancer cells transdifferentiate into contractile myCAFs with increased deposition of ECM, especially collagen, and an increased capability of tumorigenic remodeling and aligning of the ECM fibers. In cancer cells, increased stiffness of the surrounding tissue and TGFβ activity induce further expression of Jagged1, leading to a tumor-promoting feed-forward loop. Jagged1 is a central modulator of TNBC tissue structure, acting upstream of TGFβ. Created in BioRender. Parikainen, M. (2025) https://BioRender.com/o4se15z

## Discussion

The Notch signaling pathway is an attractive target for breast cancer treatment due to its roles in supporting cancer stem cells, tumor heterogeneity, and chemoresistance^35–37^. However, pan-Notch inhibition has failed clinically due to severe off-target effects, such as gastrointestinal toxicity, and the inherent complexity of the pathway where different Notch receptors and ligands play distinct and sometimes opposing roles depending on the tumor context. Targeting specific Notch components requires a precise understanding of their context-dependent roles. Here, we show that the Notch ligand Jagged1 is highly expressed in TNBC where it correlates with poor prognosis. This is in line with previous reports showing higher expression of Jagged1 in aggressive breast cancer subtypes^21–23,25,26^. We demonstrate how Jagged1 drives tumor growth and invasion by inducing CSC properties, EMT, and remodeling of the ECM in TNBC. Kumar et al. showed a similar ligand-specific role for Dll1 in promoting luminal ER+ breast cancer but not TNBC^38^. In contrast, Yamamoto and colleagues described an expansion of CSCs via NF-κB-dependent induction of Jagged1 that occurred only in basal-like breast cancer^39^. These observations further strengthen the role of individual Notch ligands in distinct breast cancer subtypes.

Breast tissue goes through major alterations during the progression towards an invasive tumor. Increased deposition of fibrillar collagens and substantial remodeling of the ECM lead to increased tissue stiffness, driving tumor progression and metastasis^3–5^. TGFβ signaling is a major driver of adverse tissue remodeling in aggressive cancer and fibrosis^40^. Our data demonstrate that Jagged1 promotes ECM deposition and fiber alignment through cancer cell-fibroblast crosstalk upstream of TGFβ, thus modulating tissue structure. We further demonstrate that higher stiffness increases Jagged1 expression. TGFβ has been shown to induce Jagged1 expression^24,41,42^, pointing towards a vicious cycle where Jagged1 induces increased tissue stiffness and TGFβ activity, which then further promote Jagged1 expression and tumor progression. Interestingly, Jagged1 can also crosstalk with the TGFβ pathway independently of Notch receptors by forming a transcriptional complex containing J1ICD, SMAD3, DDX17, and TGIF2^43^. The exact mechanisms by which Jagged1 crosstalks with TGFβ in TNBC should be further elucidated.

CAFs are the major regulators of ECM remodeling in tumors. CAFs vary in phenotype, with matrix-producing myCAFs, typically found in close proximity to cancer cells, expressing high αSMA and other TGFβ responsive genes^44^. Our data indicate that Jagged1-Notch signaling between cancer cells and fibroblasts drives myCAF activation, marked by increased αSMA expression, TGFβ signaling, and matrix production. Previous data show that αSMA is a direct Notch target gene region^45^, and other studies confirm Notch-dependent αSMA induction through cell-cell contact^13,16,46–48^. Pelon and colleagues identified four subpopulations of breast cancer-associated fibroblasts, with the subset CAF-S4 displaying a significant upregulation of Notch receptors 1-3 and demonstrating matrix remodeling and metastasis-promoting functionality^15^. The same group later identified five subpopulations of myCAFs, with TGFβ-myCAFs distinguished by a high TGFβ gene signature^31^. Interestingly, direct contact with cancer cells but not cancer cell-conditioned medium induced TGFβ-myCAF differentiation, while *NOTCH3* and *HES4* were reported among the marker genes of this fibroblast subpopulation, suggesting Notch dependency. Our data shows that Jagged1 induces differentiation of fibroblasts with a TGFβ signature and a highly similar set of marker genes as TGFβ-myCAFs, including *NOTCH3* and *HES4*. Importantly, higher abundance of TGFβ-myCAFs predicted progression of ductal carcinoma *in situ* into invasive breast cancer^49^, making them an attractive therapy target.

Previous data support the reversibility of the myofibroblastic activation state of CAFs, making CAF targeting an exciting treatment opportunity^50–52^. Targeting the stromal cells by TGFβ blockade or direct targeting of CAF markers has been shown to inhibit tumor growth and trigger anti-tumor immune responses^53–55^. Our results indicate that Jagged1 is important in mediating CAF activation, TGFβ signaling, and the development of tumor fibrosis in TNBC and suggest that therapeutically targeting Jagged1 could be an interesting avenue to control CAF phenotype and adverse TME remodeling. Of note, targeting Jagged1 might prove to be beneficial only against the more aggressive breast cancer subtypes, as we have shown it is only in these subtypes that Jagged1 expression level has a prognostic value for patient survival. In accordance with our data, a TGFβ-dependent formation of a reactive stroma through Jagged1 overexpression in cancer cells was reported in a Pten null mouse model of prostate cancer^56^, potentially extending the scope of disease contexts that might benefit from Jagged1-targeted therapy. Possible challenges and unanswered questions with this approach include whether an already formed ECM structure is reversible and whether excessive reversal of the fibrotic ECM state will lead to tumor collapse, hampering drug delivery and immune infiltration and releasing matrix-embedded, growth factors and cytokines. Future studies will need to focus on a more comprehensive analysis of Jagged1-mediated effects on other cell types of the TME besides fibroblasts, such as immune cells and adipocytes. Due to the pleiotropic effects and context-dependency of Notch signaling in the TME, the correct target and timing of Notch-based therapeutics must also be carefully planned.

## Methods

### Cell culture and treatments

MDA-MB-231, MDA-MB-436, SK-BR-3, MEF, and TIF (a kind gift from Dr. Johanna Ivaska, University of Turku, Turku, Finland) cells were grown in DMEM high glucose, supplemented with 10 % FBS, 2 mM glutamine, penicillin (100 units/mL), and streptomycin (100 µg/mL). MDA-MB-361 cells were supplemented with 20 % FBS. BT474, HCC1954, and T47D cells were grown in RPMI instead of DMEM, with same supplements. MCF10A cells were grown in DMEM/F12, supplemented with 5 % HS, 2 mM glutamine, penicillin (100 units/mL), streptomycin (100 µg/mL), 20 ng/mL EGF, 0.5 µg/mL hydrocortisone, 100 ng/mL cholera toxin, and 10 µg/mL insulin. All cells were maintained at 37 °C with 5 % CO_2_ and regularly screened against mycoplasma infections. 10 µM TGFβ receptor kinase inhibitor SB-431542 (MedChemExpress), 10 µM gamma-secretase inhibitor PF-03084014 (MedChemExpress), 500 µM LOX inhibitor β-aminopropionitrile (BAPN; MedChemExpress), or 10 ng/mL recombinant human TGFβ1 (PeproTech) were added to the medium where stated. For siRNA-mediated Jagged1 silencing, a cocktail of two siRNAs with the following sequences were used at a final concentration of 25 nM: 5’-GGG AUU UGG UUA AUG GUU A(dTdT)-3’ and 5’-GAA CCA CAG CAA CGA TCA CAA(dTdT)-3’. Non-targeting control siRNA sequence was 5’-CCU ACA UCC CGA UCG AUG AUG(dTdT)-3’. siRNAs were transfected with the Lipofectamine RNAiMAX transfection reagent (Invitrogen) according to the manufacturer’s instructions.

### Generation of Jagged1 knockout and Jagged1 rescue cells

MDA-MB-231 WT cells have endogenous expression of Jagged1. To create a Jagged1 negative cell line, sgRNA 5’-TATCAGTCCCGCGTCACGGC-3’, targeting Jagged1 exon 2, was cloned into pSPCas9(BB)-2A-GFP (PX458) (Addgene #48138)^57^ and the construct was transfected into Jag1WT cells. Transfected cells were sorted with flow cytometry as single cells on a 96-well plate based on GFP expression, allowed to grow colonies, and checked for successful Jagged1 knockout with DNA sequencing and Western blotting. 15 single cell clonal Jag1KO populations were mixed together in equal ratios to create the final Jag1KO cell line. Jag1KO cells were then transfected either with a pcDNA3.1 vector carrying Jagged1 cDNA to generate the Jag1KO+rescue cell line or with empty pcDNA3.1 vector to create the negative control cell line Jag1KO+ctrl. Jag1KO+rescue and Jag1KO+ctrl cells were grown in the presence of 500 µg/mL G418 selection.

### Chick chorioallantoic membrane (CAM) model

For monocultures, 1 x 10^6^ MDA-MB-231 cells were transplanted onto the CAM of fertilized chicken eggs on embryonal development day 8, in 20 µL of 4 mg/mL Matrigel (Corning #356231), and allowed to form tumors for 5 days, after which tumors were excised from the CAM and weighed. For co-cultures, 1 x 10^6^ MDA-MB-231 cells and 1 x 10^6^ MEF cells were used. O.C.T.-embedded co-culture CAM tumors were snap-frozen with liquid nitrogen and stored in -80 °C until further processing.

### Zebrafish invasion assay

The zebrafish used in experiments were bred and handled in the Zebrafish Core of Turku Bioscience Centre under license MMM/465/712-93 (Ministry of Agriculture and Forestry in Finland). The xenotransplantation of zebrafish embryos was performed as described in^58^. MDA-MB-231 cells were stained with 10 µM CellTracker Green CMFDA dye (Thermo Fisher Scientific) prior to xenotransplantation. 2.3 nL of cell suspension in PBS, containing 200 cells, was microinjected into the duct of Cuvier/common cardinal vein of 2 dpf zebrafish pigmentless casper strain (*roy* -/-; *mitfa* -/-) embryos. The following day, embryos were imaged with Nikon Eclipse Ti2-E microscope, with a Nikon Plan-Apochromat 2x/0.06 objective, and invaded cells were counted with ImageJ/FIJI software.

### Kaplan-Meier survival plots

Relapse-free survival of breast cancer patients was assessed in the Kaplan-Meier Plotter database for breast cancer^59^. Jagged1 216268_s_at probe set was used and patients split by auto select best cutoff. StGallen molecular breast cancer subtypes were used to visualize subtype-specific survival, and ER array status was used to split patients into ER+ and ER-subgroups.

### Single-cell data processing

The scRNA-seq data generated by Bassez et al.^27^ was obtained from https://lambrechtslab.sites.vib.be/en/single-cell. The available filtered count matrices and cell annotations were used as a starting point and reprocessed using the R package Seurat v5^60^. The code used in processing the data is provided with this paper. Briefly, treatment-naive cohort 1 samples were retrieved, low quality cells were filtered based on the QC distribution for each cell type, and data log-normalized and scaled with default parameters. Expression value 0.5 for *JAG1* was used to split TNBC cells into Jag1 high and Jag1 low populations. For fibroblasts, clustering resolution of 0.5 was used, and small subclusters of endothelial and T cells were removed based on cell type marker expression. Patients with at least 50 cancer cells in the tumor sample were retained for analysis of fibroblasts to reliably assess tumor Jag1 status. The gene set enrichment analyses were conducted with Metascape^61^, using the cell type specific transcriptomes in the dataset as background genes.

### 3D Matrigel cultures

Cell culture plates were first coated with a thin layer of 4 mg/mL Matrigel (Corning #356231). For 96-well plates, 5 x 10^3^ cells per well were seeded in 60 µL of 2 mg/mL Matrigel. For 6-well plates, 5 x 10^4^ cells per well were seeded in 600 µL of 2 mg/mL Matrigel. siRNA transfections were performed 7 hours prior to cell seeding. Matrigel was allowed to solidify o/n, after which pre-warmed cell culture medium was added on top of the 3D cultures. Spheroids were allowed to grow for 7 days. Fresh medium was changed once on day 4. Gamma-secretase inhibitor PF-03084014 containing medium was changed every 24 hours. Spheroids were extracted from Matrigel for further analysis with 5 mM EDTA in PBS. Briefly, 3D cultures were washed three times with ice-cold PBS before adding ice-cold EDTA/PBS and transferring the cultures to falcon tubes on ice. Matrigel was allowed to dissolve for 15 min before collecting the spheroids by centrifugation.

### RNA-sequencing and transcriptomics data processing

RNA was isolated using the NucleoSpin RNA kit (MACHEREY-NAGEL #740955). The sequencing library was prepared using Illumina TruSeq Stranded mRNA HT Sample Preparation kit, and sequencing was performed using Illumina NovaSeq 6000 SP with 2 x 50 bp read length at the Finnish Functional Genomics Centre, Turku, Finland. Reads were quality checked and post-processed at the Medical Bioinformatics Centre, Turku, Finland. Briefly, reads were aligned against the human reference genome (hg38), and uniquely mapped reads were used to generate gene-wise read counts. The TMM normalization from the Bioconductor package edgeR was used to normalize the filtered gene counts between the samples. Normalized counts as CPM (counts per million) values were used as input in further analysis. Genes with more than 1 CPM expression in at least 50 % of the replicates were retained in the analysis. The raw count data were transformed to CPM offsetted by 1 and log2-transformed, and then further analyzed with the Bioconductor package ROTS to acquire fold change and false discovery rate values. The gene set enrichment analysis was conducted using Metascape^61^.

### Gene expression in breast cancer patient datasets

Breast cancer patient data was accessed and downloaded through the cBioPortal service^62–64^. Jagged1 mRNA co-expression z-scores were evaluated relative to RNA-seq diploid samples in a breast invasive carcinoma dataset (TCGA, Cell 2015)^28^, with n = 817. Simple linear regression and Spearman’s rank correlation were used for modeling the correlation between mRNA expression levels, and p-values were calculated by two-tailed unpaired t-test.

### Proteomics of spheroid samples

Spheroids were made using #24-35 micro-molds from 3D Petri Dish, MicroTissues, according to the manufacturer’s instructions, with 1.4 x 10^5^ of MEFs and 1.4 x 10^5^ of either Jag1WT or Jag1KO cells mixed together and added into one mold. Medium was supplemented with 50 µg/mL ascorbic acid and changed daily. After 15 days of culture, spheroids were collected, samples processed and analyzed by liquid chromatography-mass spectrometry as described in^65^. DIA files were processed with Spectronaut (version 16.3, Biognosys) as described in^65^, using human UniProt-reviewed sequences along with their isoforms (release 2022_01) as reference. A maximum of five variable modifications per peptide was allowed. A minimum peptide length of 7 amino acids (AAs) with a maximum peptide length of 52 AAs containing a maximum of two missed cleavages was allowed. The statistical analysis was carried out using Perseus (version 1.6.14.0) based on the customized pivot report containing the MS2 quantification data from Spectronaut. The MS2 quantities were log2 transformed and normalized with width adjustment. Welch’s T-test with Benjamini-Hochberg FDR was used. ECM proteins were identified using the MatrisomeDB database (https://matrisomedb.org/)^66^.

### Immunofluorescence staining of spheroid samples

For staining of co-culture spheroids, MDA-MB-231 and MEF cells were stained with 10 µM CellTracker Orange CMRA and CellTracker Deep Red dyes (Thermo Fisher), respectively, prior to mixing cells together. Spheroids were grown the same way as mentioned previously in the proteomics section. After 15 days of culture, spheroids were collected by centrifugation, washed with PBS, and fixed and permeabilized in PBS containing 4 % PFA and 1 % Triton X-100 for 3 h at +4 °C. After fixation, spheroids were washed in PBST (0.1 % Triton X-100 in PBS) for 15 min and then incubated in blocking buffer (3 % BSA in PBST) for 3 h at +4 °C. Spheroids were washed for 15 min in PBST and then incubated in 20 ng/µL CNA35 (pET28a-EGFP-CNA35 was a gift from Maarten Merkx; Addgene plasmid #61603^32^) and 600 nM DAPI in PBST overnight at +4 °C with gentle shaking. Spheroids were washed three times in PBST and mounted on microscopy slides in glycerol. A Zeiss LSM880 Axio Observer.Z1 confocal microscope and ZEN 2.3 SP1 black edition software were used for imaging. The objective used was a Zeiss Plan-Apochromat 20x/0.8.

### Western blotting

Spheroids were lysed on ice with Laemmli SDS sample buffer containing 3 % β-mercaptoethanol and protein lysates denatured by boiling. Proteins were separated by SDS-PAGE and transferred to a nitrocellulose membrane using a wet transfer apparatus, followed by blocking with 5 % nonfat dry milk in TBS. Membranes were incubated with primary antibodies diluted in 3 % BSA/0.02 % NaN_3_ overnight at +4 °C, followed by incubation with a secondary antibody, coupled to horseradish peroxidase, for 1 h RT. The following antibodies and dilutions were used: 1:1000 Jagged1 (Cell Signaling Technology #28H8), 1:1000 phospho-Smad2 (Ser465/467)/Smad3 (Ser423/425) (Cell Signaling Technology #8828), 1:1000 Smad2/3 (Santa Cruz #sc-133098), 1:1000 αSMA (Cell Signaling Technology #19245), 1:2500 FN (BD Pharmingen #610077), 1:1000 ERα (Santa Cruz #sc-8002), 1:1000 β-tubulin (Cell Signaling Technology #86298), 1:200 000 β-actin (Sigma #A1978), and 1:5000 HSC70 (Enzo Life Sciences #ADI-SPA-815). Proteins were detected with SuperSignal™ West Pico PLUS Chemiluminescent Substrate (Thermo Scientific) using the iBright FL1500 Imaging System (Invitrogen). The results were normalized against housekeeping protein expression and by using the sum of all data points method^67^. For collagen visualization, dot blot technique was used. Cell lysates were directly spotted on a nitrocellulose membrane, allowed to dry, rinsed with MQ-H_2_O, and stained for total protein using the Revert™ 700 Total Protein Stain Kit (Li-COR) according to manufacturer’s instructions. After destaining, nonspecific binding was blocked with 5 % nonfat dry milk in TBS and the membrane incubated with 20 ng/µL CNA35 in PBS for 1 h RT. Membrane was washed with TBS and GFP signal imaged using the iBright FL1500 Imaging System (Invitrogen). ImageJ/FIJI software was used to quantify images.

### Cell-derived matrices

CDMs were produced as previously described in^68^. Briefly, 6 x 10^5^ fibroblast cells for monocultures or 2 x 10^5^ MDA-MB-231 and 4 x 10^5^ fibroblast cells for co-cultures were plated on gelatin-coated coverslips and allowed to form a confluent monolayer overnight. The following day, treatment with 50 μg/mL ascorbic acid was started and continued for 12 or 15 days for TIF or MEF co-cultures, respectively. Ascorbic acid containing medium was changed daily. After cell denudation, CDMs were fixed and stained as described in^68^. For fibronectin staining, 2.5 μg/mL mouse anti-fibronectin antibody (BD Pharmingen #610077) was used, followed with 4 μg/mL anti-mouse Alexa Fluor 555-conjugated secondary antibody (Invitrogen #A-21424). Collagen was visualized with 20 ng/µL CNA35, incubated together with the secondary antibody. CDMs were imaged using a Zeiss LSM880 Axio Observer.Z1 confocal microscope using ZEN 2.3 SP1 black edition software. The objective used was a Zeiss Plan-Apochromat 20x/0.8. One z slice from the middle of the sample was captured, and five images per sample at randomized positions were acquired for analysis. Fiber alignment coefficients were analyzed with the CurveAlign V4.0 Beta software^69,70^. Fibers in image boundary were excluded from analysis and 3^rd^ finest scale used.

### Recombinant Notch ligand coating

Gelatin-coated coverslips were incubated with 50 µg/mL recombinant Protein G (Thermo Fisher) in PBS at RT overnight. Coverslips were washed three times with PBS, blocked with 1 % BSA/PBS for 1 h RT, and washed three times with PBS. Coverslips were then incubated with 10 nM recombinant human Fc-linked Notch ligands (R&D Systems; Jag1 #1277-JG, Dll1 #10184-DL, Dll4 #10185-D4) or 10 nM recombinant human Fc-linked IgG1 (R&D Systems #110-HG) in 0.1 % BSA/PBS overnight at +4 °C. Coverslips were washed three times with PBS before plating cells for CDM production.

### CAM cryosection staining

Snap-frozen CAM tumors in O.C.T. were cut into 8 µm cryosections for immunofluorescence staining and 6 µm cryosections for hematoxylin and eosin staining. For immunofluorescence staining, cryosections were fixed and permeabilized with 4 % PFA/1 % Triton X-100 in PBS for 10 min RT, washed with PBS, and incubated in blocking buffer (10 % horse serum in PBS) for 30 min RT. Primary antibodies against fibronectin (BD Pharmingen #610077), αSMA (Cell Signaling Technology #19245), and pan-cytokeratin (Sigma #C2562) were diluted 1:100 in blocking buffer and incubated on cryosections overnight at +4 °C. The following day, cryosections were washed three times with PBST and incubated with 4 μg/mL secondary antibodies (anti-mouse Alexa Fluor 555, Invitrogen #A-21424, and anti-rabbit Alexa Fluor 647, Invitrogen #A-31573), 165 nM Alexa Fluor 633 Phalloidin (Invitrogen #A22284), 20 ng/µL CNA35, and 600 nM DAPI in blocking buffer for 1 h RT. Cryosections were washed three times with PBS, rinsed once with MQ-H_2_O, and mounted with Mowiol/DABCO on microscopy slides. A Zeiss LSM880 Axio Observer.Z1 confocal microscope and ZEN 2.3 SP1 black edition software were used for imaging, with Zeiss EC Plan-Neofluar 10x/0.30 objective. Integrated signal densities were quantified using the ImageJ/FIJI software.

### Quantitative PCR

RNA was isolated using the NucleoSpin RNA kit (#740955, MACHEREY-NAGEL) and cDNA was prepared from 1 µg of RNA using the SensiFAST cDNA Synthesis kit (Bioline). The reaction mixtures for quantitative PCR were prepared with 5xHOT FIREPol EvaGreen qPCR Mix Plus with ROX (Solis BioDyne) with 10 µL final reaction volume, containing 4 µL of cDNA and primers at a 250 nM final concentration. Primer sequences are provided in Supplementary Table 6. C_T_ values were normalized to UBC C_T_ values, and fold changes were calculated with the 2^-ΔΔCt^ formula

### Protein expression on increasing substrate stiffness

MDA-MB-231 or MCF7 WT cells were plated on Matrigen Softwell Petrisoft 35 mm Dish Collagen (Cell Guidance Systems) dishes with 0.5, 4, or 50 kPa stiffness for 72 h before cell lysis with Laemmli SDS sample buffer containing 3 % β-mercaptoethanol. Lysates were analyzed by Western blotting as described above.

### Statistics and reproducibility

Statistical analyses and illustrations were prepared using GraphPad Prism 10.3.1 or R/R-Studio (v4.4.2). Experiments were independently repeated a minimum of three times. Detailed information about exact *n* values, displayed data and error bars, and statistical tests are reported in the figure legends. Normality of the data was tested using the Shapiro-Wilk test prior to the use of any parametric tests.

## Acknowledgements

Alexandra Manea and Kai-Lan Lin are thanked for their technical assistance. Dr. Valeriy Paramonov is acknowledged for designing the Jagged1 CRISPR-Cas9 construct. We acknowledge the Finnish Functional Genomics Centre, the Cell Imaging and Cytometry Core, and the Zebrafish Core at Turku Bioscience Centre, all supported by Biocenter Finland, for their excellent service. We thank the Medical Bioinformatics Centre of the Turku Bioscience Centre for the RNA-sequencing data analysis. The Centre is supported by the University of Turku, Åbo Akademi University, Biocenter Finland, and ELIXIR Finland. This project has received funding from the European Union’s Horizon 2020 research and innovation program under grant agreement #953234 (Tumor-LN-oC; C.S.) and the Academy of Finland under decision numbers #307133 (C.S., M.P.) and #309373 (PESCADoR; C.S.). This research was supported by the Research Council of Finland’s Flagship InFLAMES (decision numbers #337531 and #357911). The Åbo Akademi University Foundation’s Center of Excellence in Cellular Mechanostasis (CellMech, C.S.), Cancer Foundation Finland sr (C.S., M.P.), the Jane and Aatos Erkko Foundation (C.S., M.P.), and the Sigrid Jusélius Foundation (C.S., M.P.) have also supported the research. Additionally, M.P. has received funding from the Varsinais-Suomi Regional Fund of the Finnish Cultural Foundation, the Swedish Cultural Foundation in Finland, the K. Albin Johansson Foundation, and the Ida Montin Foundation.

## Author contributions

M.P., J.H., and C.S. designed the experiments. U.S. and P.R. conducted the mass spectrometry analysis and related data analysis of proteomics samples. M.P. performed the rest of the experiments, analyzed data, performed statistical analyses, and prepared the figures. M.P., J.H., and C.S. wrote the manuscript with input from all the authors.

## Competing interests

The authors declare no competing interests.

---

**Extended Data Figure 1.**
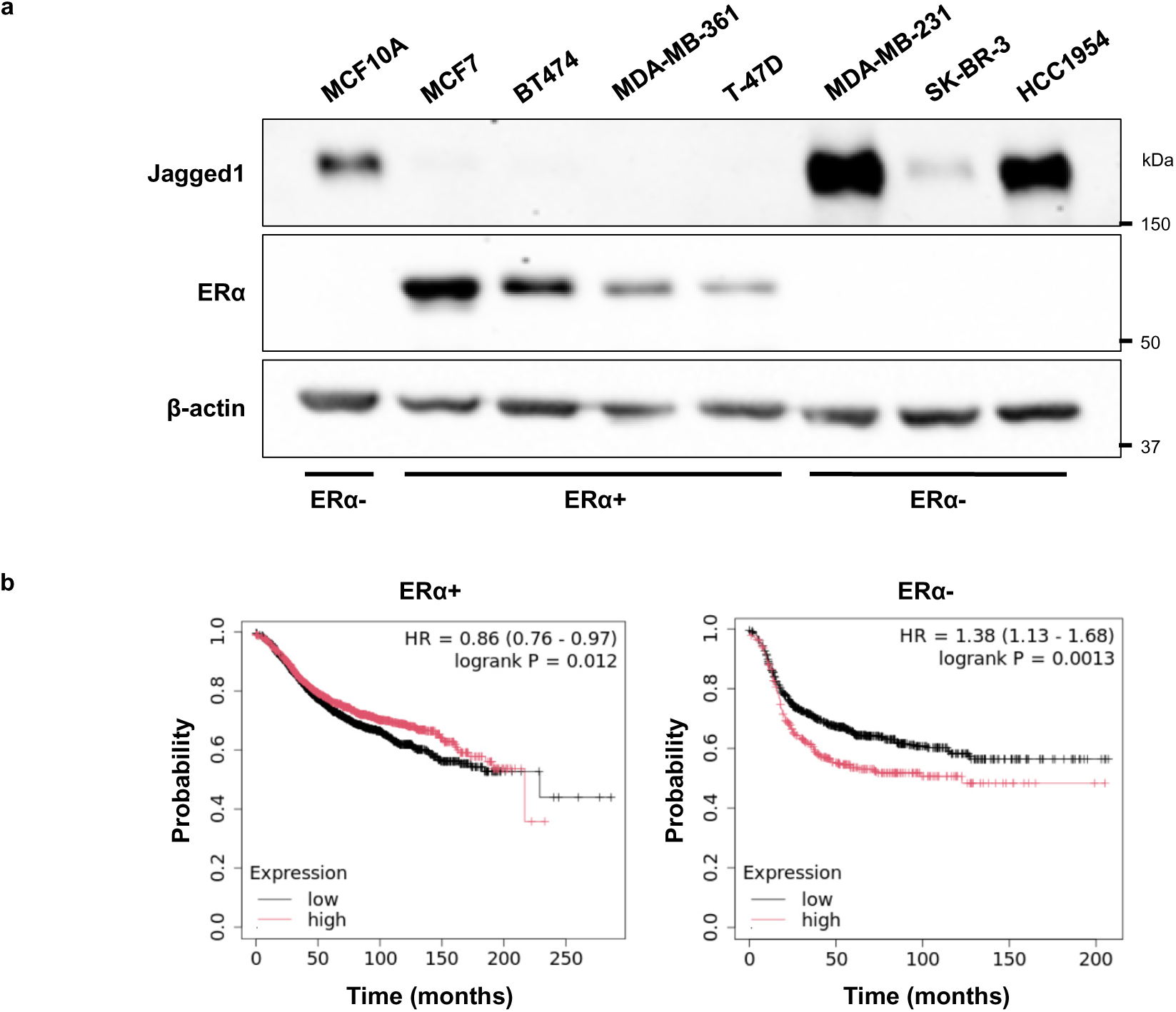
High Jagged1 expression is associated with estrogen receptor (ERα) negative breast cancer. **a** Western blots showing Jagged1 and ERα expression levels in a panel of breast cancer cell lines and in the non-tumorigenic, ERα negative breast epithelial cell line MCF10A. **b** Relapse-free survival of patients with ERα+ or ERα-breast cancer and either low (black) or high (red) expression of Jagged1.

**Extended Data Figure 2.**
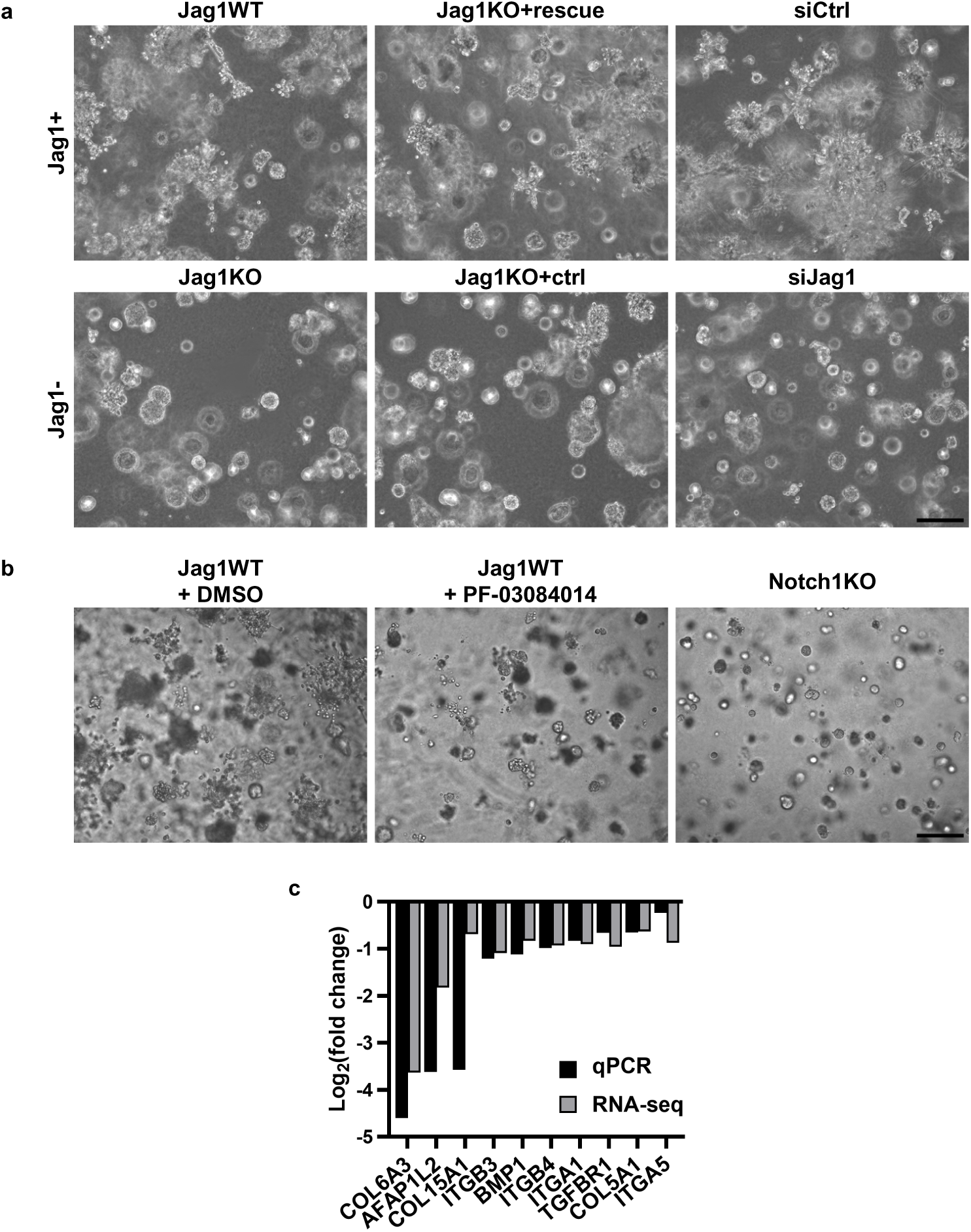
Modulation of Jagged1-Notch signaling affects breast cancer spheroid growth and invasion. Representative spheroid images of **a** MDA-MB-231 Jag1WT and Jag1KO cells, Jag1KO cells transfected with Jag1-containing plasmid (Jag1KO+rescue) or empty vector plasmid (Jag1KO+ctrl), and Jag1WT cells transfected with non-targeting siRNA (siCtrl) or Jag1-targeting siRNA (siJag1), and **b** Jag1WT cells with inhibited Notch signaling activity either by a gamma-secretase inhibitor (PF-03084014) or knockout of Notch1 (Notch1KO). Scale bar 200 µm. **c** Log2 fold changes of genes downregulated in Jag1KO cells, determined by quantitative PCR (qPCR) or whole transcriptome sequencing (RNA-seq). Data are presented as mean of three independent experiments.

**Extended Data Figure 3.**
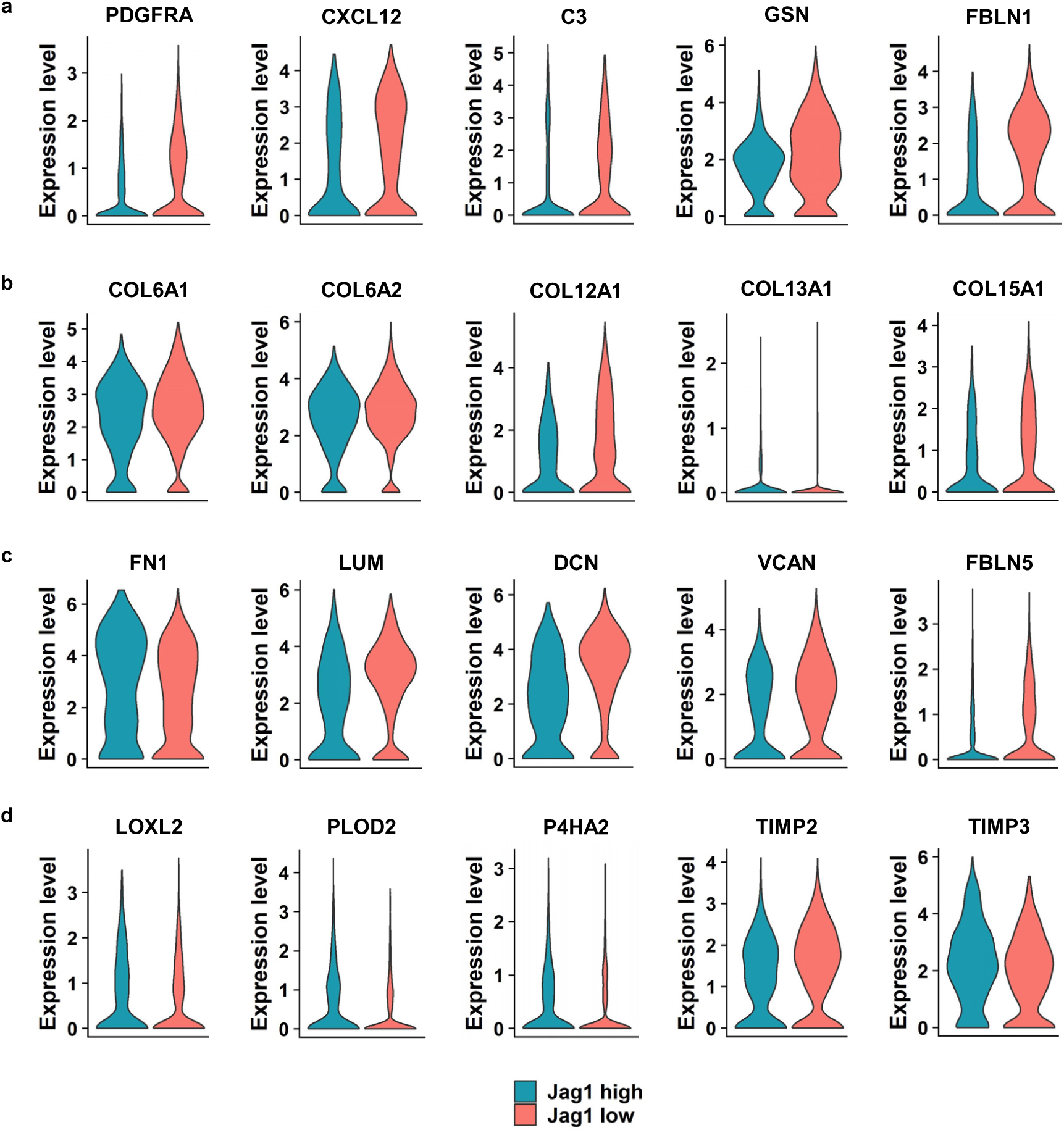
Jagged1 expressed by cancer cells regulates expression of ECM and ECM remodeling genes in fibroblasts of the same tumors. Expression of **a** iCAF markers, **b** collagens, **c** ECM genes, and d ECM remodeling enzymes in scRNA-seq data of fibroblasts from TNBC tumors with either high or low expression of Jagged1 in cancer cells. n = 881 of Jag1 high fibroblasts and n = 1530 of Jag1 low fibroblasts.

**Extended Data Figure 4.**
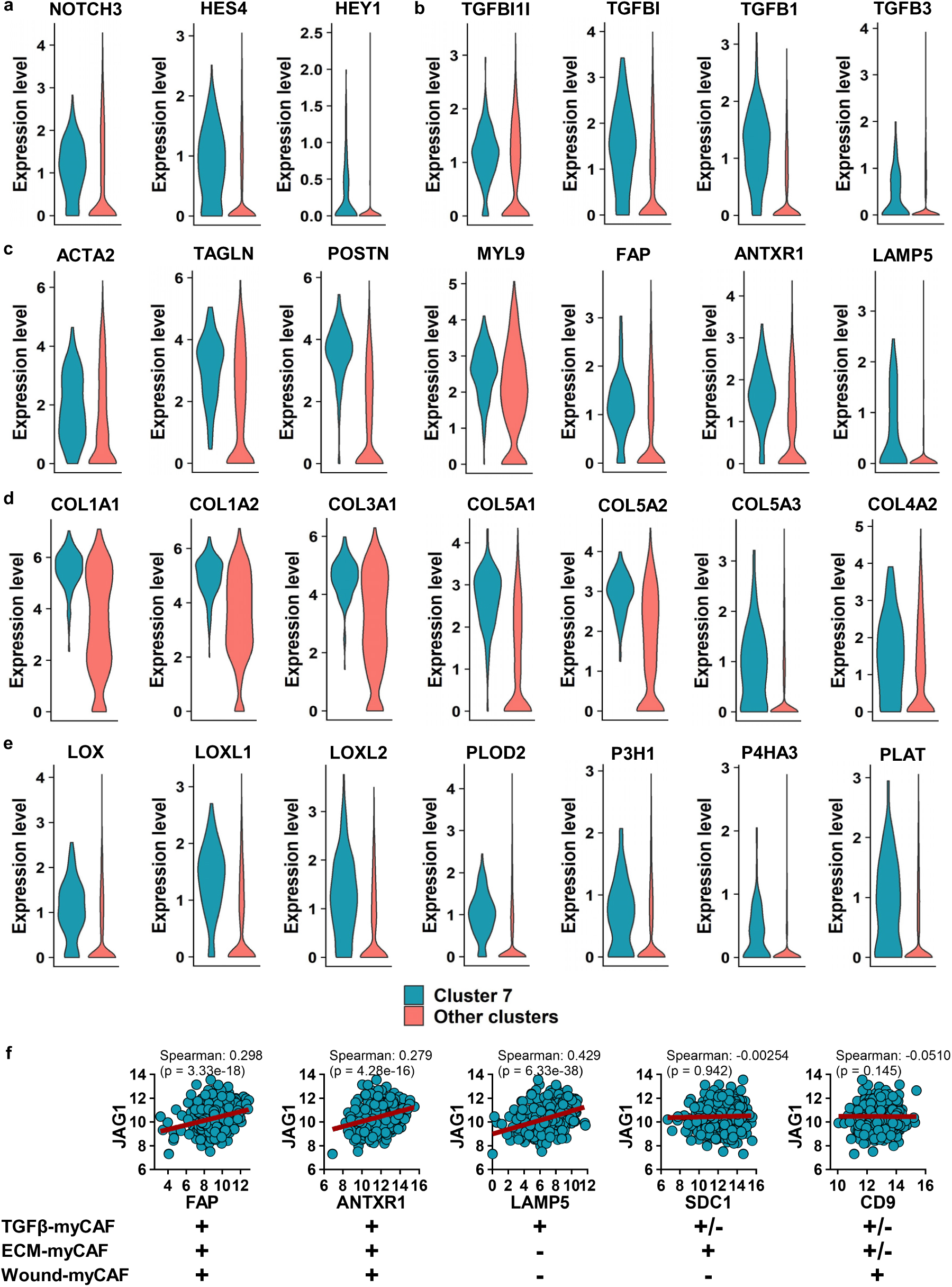
High Jagged1 expression in cancer cells promotes differentiation of fibroblasts into a specific subtype marked by high Notch, TGFβ, and myCAF gene signatures. Expression of **a** Notch, **b** TGFβ, **c** CAF marker, **d** collagen, and **e** ECM remodeling enzyme genes in fibroblast cluster 7 in scRNA-seq data of TNBC tumors. n = 176 of cluster 7 and n = 2235 of other cluster fibroblasts. **f** Correlation of *JAG1* expression with markers of different myCAF subpopulations in a breast cancer patient mRNA expression dataset (TCGA, Cell 2015, n = 817).

**Extended Data Figure 5.**
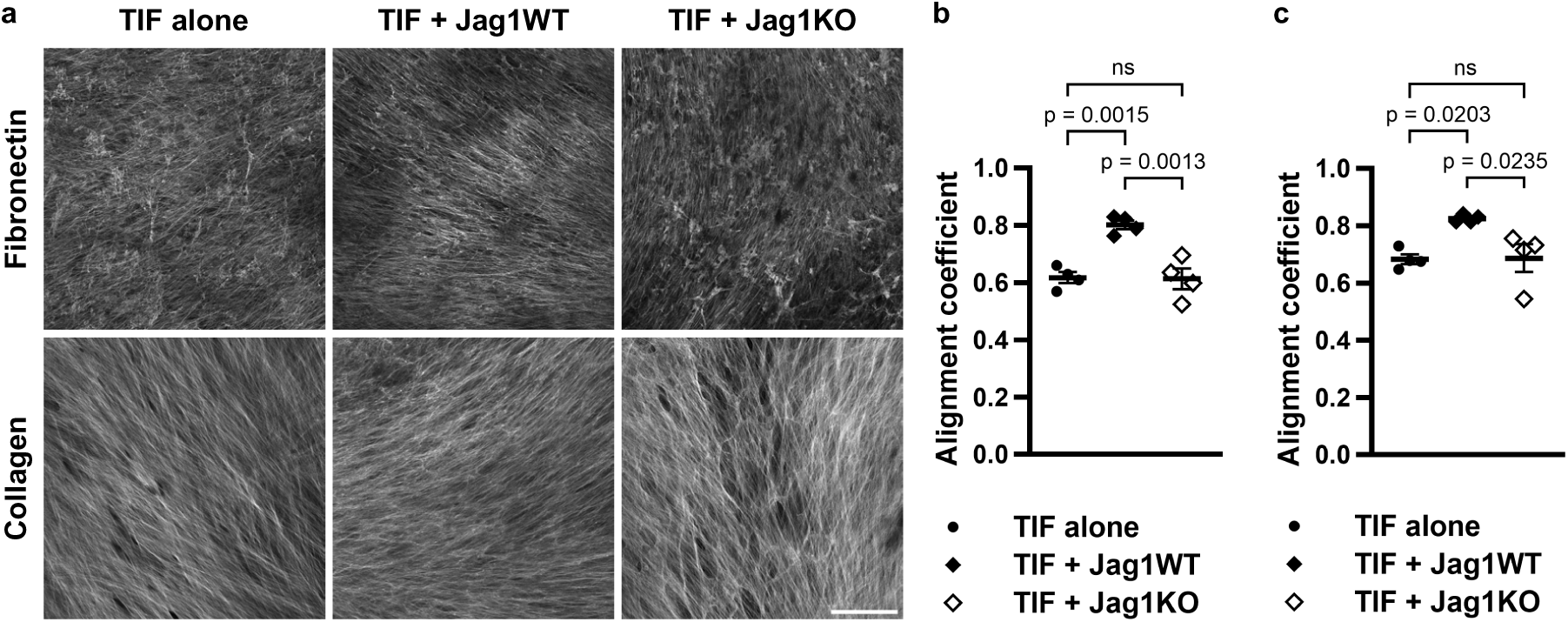
Jagged1 expressed by cancer cells directs extracellular matrix fiber alignment in co-culture with human fibroblasts. **a** Representative images of fibronectin and collagen stainings of cell-derived matrices from human telomerase immortalized fibroblast (TIF) monocultures or TIF co-cultures with MDA-MB-231 Jag1WT or Jag1KO cells, and alignment coefficients of **b** fibronectin and c collagen fibers. Data are presented as mean ± SEM of four independent experiments. p-values are calculated by one-way ANOVA with Tukey’s multiple comparisons test. Scale bar 100 µm.

**Extended Data Figure 6.**
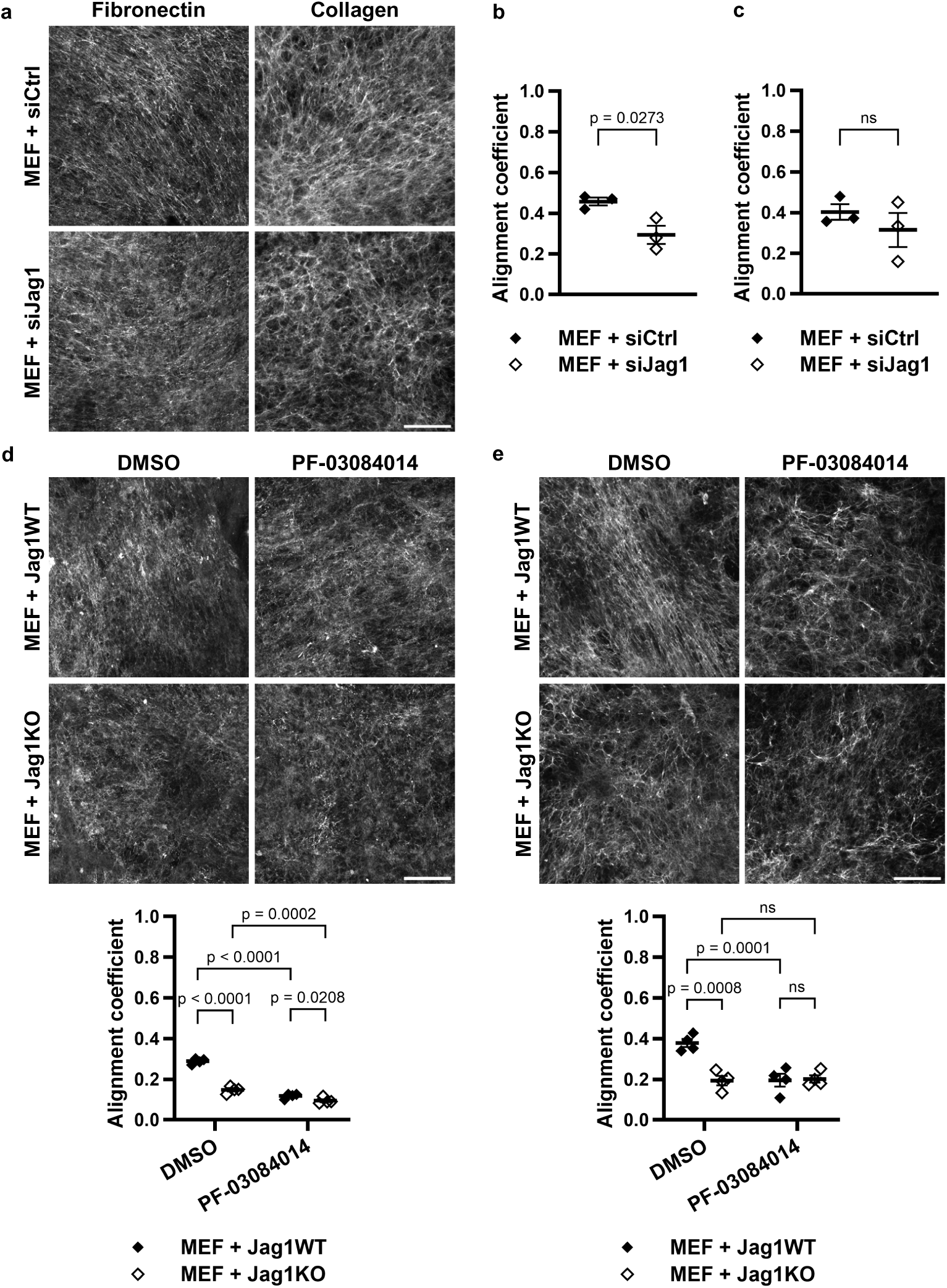
Jagged1 expressed by cancer cells directs extracellular matrix fiber alignment via activation of Notch signaling. **a** Representative images of fibronectin and collagen stainings of cell-derived matrices (CDM) from MEF co-cultures with MDA-MB-436 cells transfected with either non-targeting siRNA (siCtrl) or Jag1-targeting siRNA (siJag1), and alignment coefficients of **b** fibronectin and **c** collagen fibers. p-values are calculated by two-tailed unpaired t-test. Data are presented as mean ± SEM of three independent experiments. Representative images and fiber alignment coefficients of **d** fibronectin and **e** collagen of CDMs from DMSO or PF-03084014 gamma-secretase inhibitor treated MDA-MB-231 Jag1WT or Jag1KO co-cultures with MEF cells. p-values are calculated by two-way ANOVA. Data are presented as mean ± SEM of four independent experiments. Scale bar 100 µm.

**Extended Data Figure 7.**
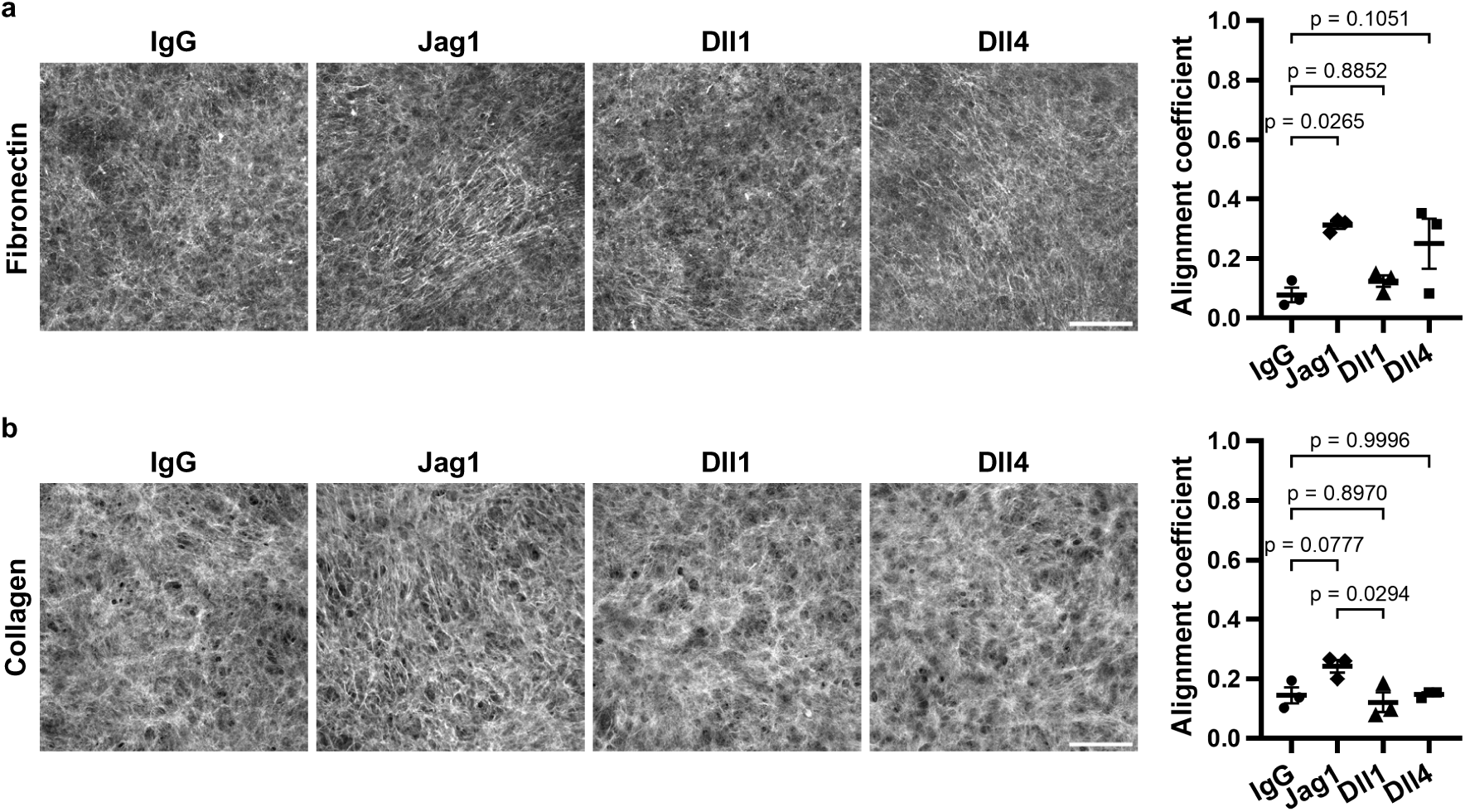
Recombinant Jagged1 directs extracellular matrix fiber alignment. Representative images and fiber alignment coefficients for **a** fibronectin and **b** collagen from cell-derived matrices of MEF cells cultured on recombinant IgG, Jag1, Dll1, or Dll4 coated coverslips. p-values are calculated by one-way ANOVA with Tukey’s multiple comparisons test. Data are presented as mean ± SEM of three independent experiments. Scale bar 100 µm.

**Extended Data Figure 8.**
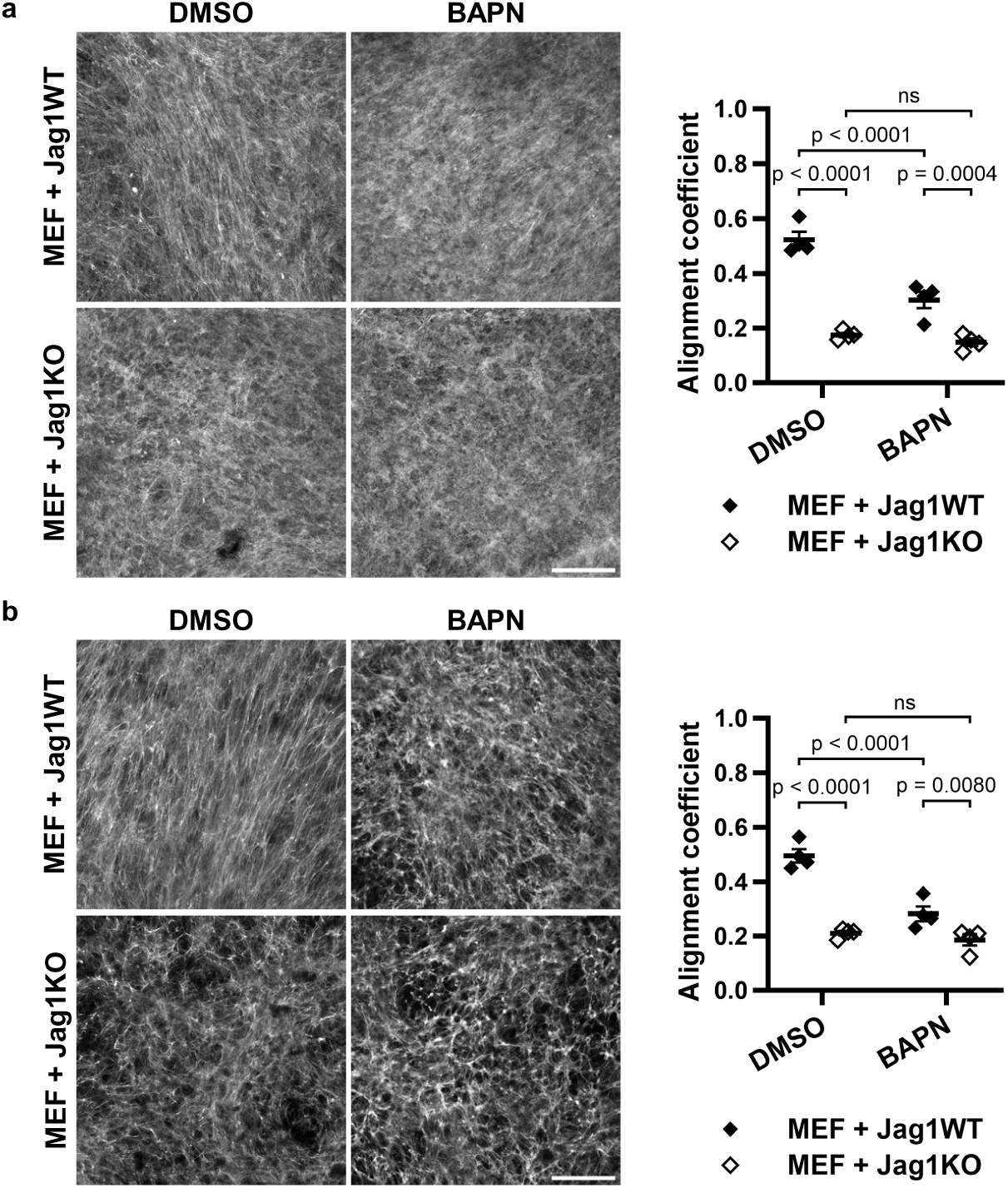
Lysyl oxidase inhibition partly abrogates the induction of extracellular matrix fiber alignment by Jagged1. Representative images and fiber alignment coefficients for a fibronectin and b collagen from cell-derived matrices of MEF co-cultures with MDA-MB-231 Jag1WT or Jag1KO cells treated with either DMSO or BAPN (β-aminopropionitrile), a lysyl oxidase inhibitor. p-values are calculated by two-way ANOVA. Data are presented as mean ± SEM of four independent experiments. Scale bar 100 µm.

**Extended Data Figure 9.**
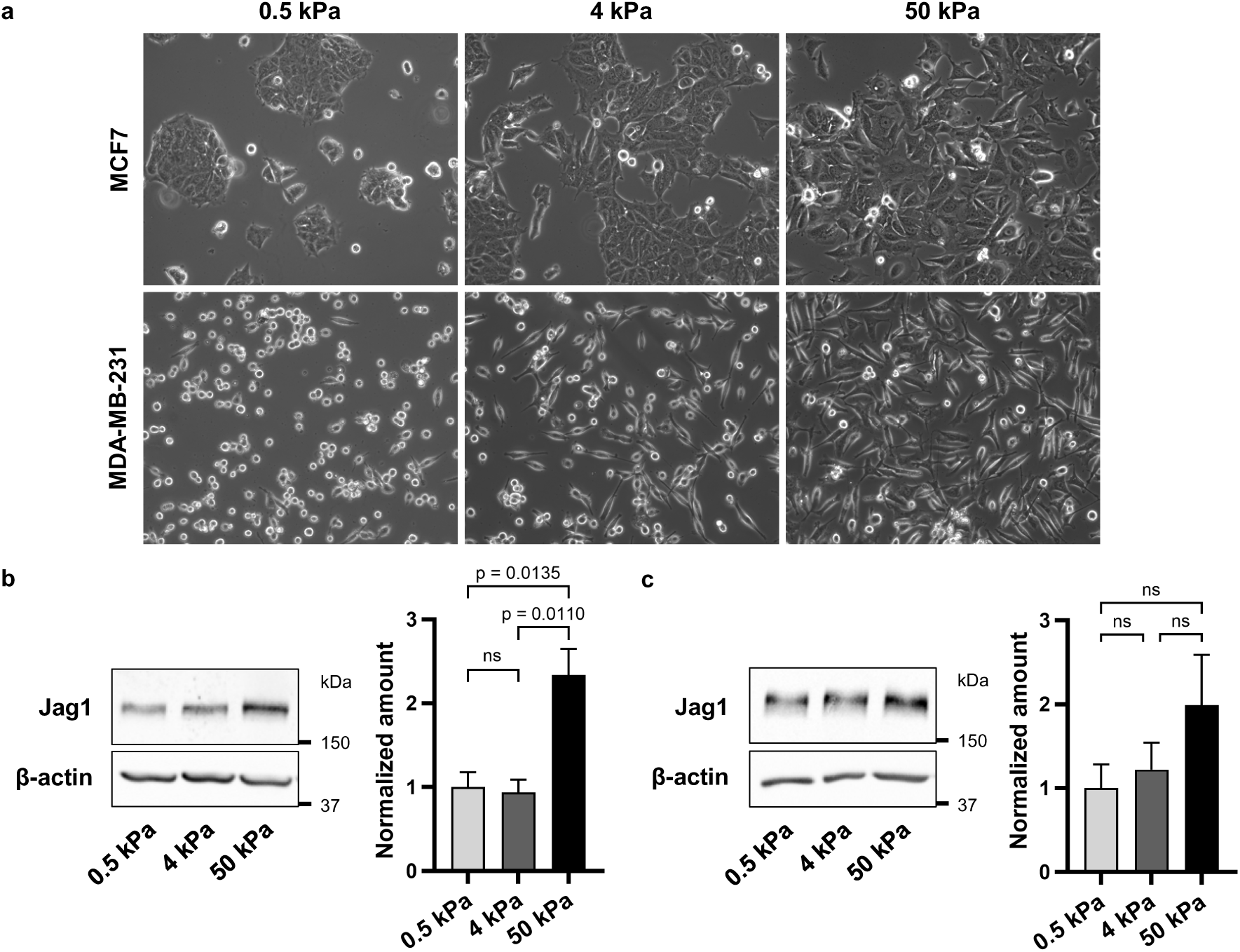
Matrix stiffness promotes Jagged1 protein expression. **a** Representative images of MCF7 and MDA-MB-231 wild-type cells grown on increasing substrate stiffness. Western blot analysis of Jag1 levels in **b** MCF7 and **c** MDA-MB-231 cells grown on increasing substrate stiffness. Quantifications of three independent experiments are shown to the right. Data are presented as mean ± SEM. p-values are calculated by one-way ANOVA with Tukey’s multiple comparisons test.

**Extended Data Figure 10.**
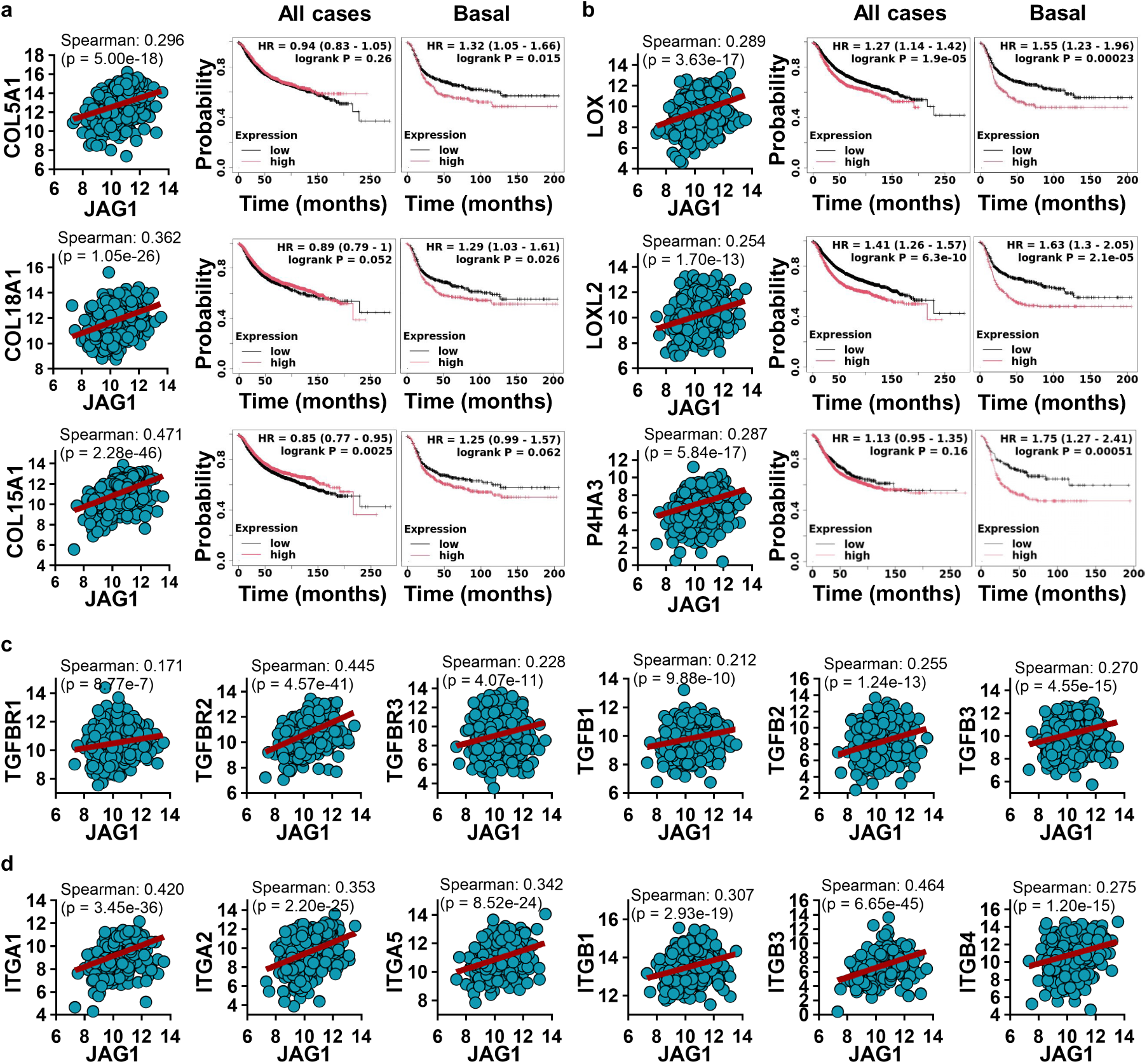
Jagged1 expression correlates with ECM, TGFβ, and integrin gene expression in breast cancer patient data. mRNA expression level correlation with Jag1 mRNA expression in breast cancer patients (TCGA, Cell 2015, n = 817) and relapse-free survival of patients with low (black) or high (red) expression of **a** collagens *COL5A1*, *COL18A1*, and *COL15A1* and **b** extracellular matrix modifying enzymes *LOX*, *LOXL2*, and *P4HA3* in all breast cancer cases and in basal breast cancer separately. mRNA expression level correlation with Jag1 mRNA expression in breast cancer patients (TCGA, Cell 2015, n = 817) of **c** TGFβ receptors and ligands *TGFBR1*, *TGFBR2*, *TGFBR3*, *TGFB1*, *TGFB2*, and *TGFB3* and **d** integrins *ITGA1*, *ITGA2*, *ITGA5*, *ITGB1*, *ITGB3*, and *ITGB4*.

## Notes

### Competing Interest Statement

The authors have declared no competing interest.

